# The vestibular calyceal junction is dismantled following subchronic streptomycin in rats and sensory epithelium stress in humans

**DOI:** 10.1101/2022.05.17.492294

**Authors:** Alberto F. Maroto, Mireia Borrajo, Sílvia Prades, Àngela Callejo, Emilio Amilibia, Marta Pérez-Grau, Francesc Roca-Ribas, Elisabeth Castellanos, Alejandro Barrallo-Gimeno, Jordi Llorens

## Abstract

Hair cell (HC) loss by epithelial extrusion has been described to occur in the rodent vestibular system during chronic 3,3’-iminodipropionitrile (IDPN) ototoxicity. This is preceded by dismantlement of the calyceal junction in the contact between type I HC (HCI) and calyx afferent terminals. Here, we evaluated whether these phenomena have wider significance. First, we studied rats receiving streptomycin for 3 to 8 weeks. Streptomycin caused loss of vestibular function associated with partial loss of HCI and decreased expression of contactin-associated protein (CASPR1), denoting calyceal junction dismantlement, in the calyces encasing the surviving HCI. Additional molecular and ultrastructural data supported the conclusion that HC-calyx detachment precede HCI loss by extrusion. Animals allowed to survive after the treatment showed functional recuperation and rebuilding of the calyceal junction. Second, we evaluated human sensory epithelia obtained during therapeutic labyrinthectomies and trans-labyrinthine tumour excisions. Some samples showed abnormal CASPR1 label strongly suggestive of calyceal junction dismantlement. Therefore, reversible dismantlement of the vestibular calyceal junction may be a common response triggered by chronic stress, including ototoxic stress, before HCI loss. This may partly explain clinical observations of reversion in function loss after aminoglycoside exposure.

## INTRODUCTION

Sensory encoding by the inner ear’s vestibular system depends on the transducing function of hair cells (HC). In most mammals, permanent loss of equilibrium and gaze control occurs as a result of HC loss considering that these cells do not regenerate to a significant extent to restore functionality. Nevertheless, significant recuperation in vestibular function is recorded clinically in a variety of disease conditions. In some cases, such as benign paroxysmal positional vertigo (von Brevern et al., 2015), recuperation can be complete, suggesting that the condition is associated with pathogenic mechanisms that transiently impair the system’s function, but do not cause HC loss. In other disease conditions, such as Ménière’s disease (López-Escámez et al., 2015), recurrent crises are often associated with incomplete recuperation and progressive worsening of the vestibular function. In the case of exposure to ototoxic aminoglycoside antibiotics, irreversible loss of vestibular function is a common outcome (Black et al., 2004), but significant yet incomplete recuperation has also been recorded (Black et al., 2001). In recent years, we have used a sub-chronic ototoxicity model to identify reversible cellular and molecular alterations in the vestibular epithelia that associate with reversible loss of vestibular function in rats (Sedó-Cabezón et al., 2015) and mice (Greguske et al., 2019). These studies showed that the sub-chronic ototoxic stress caused by the experimental compound 3,3’-iminodipropionitrile (IDPN) results in the dismantlement of the calyceal junction between type I HCs (HCI) and the calyx afferent terminals, as demonstrated by ultrastructural characterization and observed loss of the junction proteins CASPR1 (contactin-associated protein) and tenascin-C, as well as delocalization of the voltage-gated potassium channel KCNQ4. Simultaneously, the vestibular epithelium showed a loss of pre-synaptic and post-synaptic proteins, indicative of synaptic uncoupling between the HCs and their afferents. These changes that are associated with the loss of vestibular function are revealed by visible behavioural deficits. In this model, if the exposure was halted at an early stage of the toxicity, both the epithelium alterations and the behavioural dysfunction were largely reversible. In contrast, if the exposure was maintained, the functional loss persisted as a result of HC loss, which predominantly occurred by the extrusion of live cells from the sensory epithelium into the endolymphatic cavity (Llorens and Rodríguez-Farré, 1997; Seoane et al., 2001a,b; Sedó-Cabezón et al., 2015; Greguske et al., 2019).

The rat and mouse sub-chronic IDPN models illustrate a cellular pathway to HC demise that differs from apoptosis (Seoane et al., 2001a). As explained in the previous paragraph, it concludes with the extrusion of HCs after a phase of sensory cell detachment and synaptic uncoupling (Sedó-Cabezón et al., 2015; Greguske et al., 2019). The fact that extrusion is the main form of HC demise in birds and amphibians following a variety of insults (Hirose et al., 1999; Rubel et al., 2013), suggests that its molecular mechanisms may be highly conserved across vertebrates and operate in response to diverse forms of damage. Surprisingly, this phenomenon has received scarce attention and information is lacking about its molecular mechanisms and relevance to human health. One first question worth addressing is whether these phenomena have a role in mammalian aminoglycoside toxicity *in vivo*, as this is the main cause of vestibular toxicity in humans. Although most research on aminoglycoside-induced HC loss has focused on apoptosis (Furness, 2015), the literature also contains evidence that these drugs may cause vestibular HC extrusion (Li et al., 1995; Nakagawa et al., 1997; Granados and Meza, 2005). In the present work, we studied whether dismantlement of the calyceal junction, synaptic uncoupling, and HC extrusion occur in the rat vestibular epithelium in response to one aminoglycoside antibiotic, streptomycin. Previously, we found that the mode of HC demise, whether extrusion, apoptosis, or necrosis was determined by the intensity and duration of the toxic stress. Thus, high acute doses of IDPN caused HC necrosis, sub-acute exposure to medium doses caused HC apoptosis and sub-chronic low dose exposure elicited HC extrusion (Seoane et al., 2001a). These data indicate that short exposure paradigms aimed at accelerating and synchronizing HC demise (Taylor et al., 2008; Oesterle et al., 2008; Benkafadar et al., 2021) may in fact favour the apoptotic pathways above other possible pathways. Accordingly, for the present study we investigated the effects of sub-chronic aminoglycoside exposure. As a second approach to evaluate the relevance of these phenomena to human disease conditions, we explored their occurrence in the vestibular epithelia of a substantial number of patients with vestibular schwannoma and a few with other vestibular diseases that also require trans-labyrinthine tumour excision (meningioma and endolymphatic sac tumour) or therapeutic labyrinthectomy (Ménière’s disease). Vestibular schwannomas are tumours that grow in the vestibular nerve and cause hearing and vestibular function loss. Current evidence indicates that this loss may in part be caused by the negative impact pro-inflammatory factors secreted by the tumour have on the sensory epithelia (Dilwali et al., 2015). The data we obtained demonstrate that calyceal junction dismantlement occurs during chronic streptomycin toxicity in rats and provide evidence that it occurs in human vestibular epithelia in response to other types of stress.

## MATERIALS AND METHODS

### Animals and treatments

Animals were used in accordance with Law 5/1995 and Act 214/1997 of the Generalitat de Catalunya and approved by the University of Barcelona’s Ethics Committee on Animal Experiments and the Commission on Animal Experimentation of the Generalitat. Long-Evans rats were obtained from Janvier Labs (Le-Genest-Saint-Isle, France) at post-natal day 8-12. They were housed with lactating mothers in standard Macrolon cages (215×465×145 mm) with wood shavings as bedding. At day 21, the rats were weaned and housed in groups of four of the same sex. Housing conditions included a 12:12 h light:dark cycle (07:30-19:30 h), a 22±2°C room temperature and an *ad libitum* access to standard pellet diets (TEKLAD 2014, Harlan Laboratories, Sant Feliu de Codines, Spain).

A total of 98 rats (58 males, 40 females) were used in two different administration protocols, one injection per day and two injections per day. The first administration protocol was designed on the grounds of the protocols by Granados and Meza, 2005, and Schirmer et al., 2007. Animals were administered streptomycin sulphate salt (Sigma-Aldrich, S6501), s.c., dissolved in PBS, starting at postnatal 21, daily for 6 or 8 weeks. To ensure dose accuracy, injection volumes were 10 ml/kg at the beginning, but were reduced to 2 ml/kg as the rats increased body weights. Different injection sites in the dorsal skin were used in an alternating sequence. In this first series of experiments, rats were dosed with 0 (control vehicle), 100, 300, 400 or 500 mg/kg · day of streptomycin (referred to the free base), once a day, for 6 or 8 weeks. Two experimental batches used only males, one of them used only females, and the fourth one included an equal number of males and females. Acute toxicity and lethality is described in the results section above. A final experiment used a second administration protocol, in which the total daily dose was split into two injections given 8 h apart. This dosing paradigm was designed to evaluate the vestibular effects of larger daily doses and circumvent the acute toxicity of streptomycin recorded in rats receiving the 500 mg/kg dose. Thus, rats received 0, 600, 700 or 800 mg/kg · day in two daily doses of 0, 300 mg/kg (for 6 weeks), 350 mg/kg (for 4 weeks) or 400 mg/kg (for 3 weeks). No animals died in this experiment. The detail of all groups of animals used is shown in Supplementary Table S1.

During the period of treatment, the animals were weighted and assessed for wellbeing daily. They were also tested for vestibular function once a week. At the end of the treatment period, some animals were allowed to survive for up to 8-10 additional weeks to evaluate the potential reversibility of the vestibular dysfunction and the histopathological effects of streptomycin. Some control animals were also allowed to survive the recovery period to provide control body weight and vestibular function data. The day after that of the last dose, or after the recovery period, rats were euthanized by decapitation under anaesthesia. The skull was sectioned in half and placed in fixative solution under a fume hood for dissection of the vestibular sensory epithelia. The epithelia from the first ear were typically obtained within 5 min after decapitation and used for scanning (SEM) and transmission (TEM) electron microscopy. The epithelia from the second ear, obtained within 15 min after decapitation, were used for immunohistochemistry. Epithelia from some animals were used for immunohistochemical analysis only. The numbers of animals used in each histological analysis are indicated in the figures and are summarized in Suppl. Table S2.

### Human samples

Vestibular sensory epithelia were collected from patients submitted to therapeutic labyrinthectomy or translabyrinthine tumour excision. The samples included in the study, detailed in Suppl. Table S3, are from 2 cases of Ménière’s Disease, 1 case of endolymphatic sac tumour, 4 cases of meningioma, and 22 cases of vestibular schwannoma. Patients were 18 to 70 years old, and 52 % were females. The tissues were obtained after informed consent, as approved by the Clinical Ethics Committee of the Hospital Germans Trias i Pujol. The samples were immediately placed in 4% formaldehyde with <2.5% methanol (BiopSafe) and sent to the laboratory, usually in 24 h, although some samples were delayed for up to 7 days (Suppl. Table S3). The tissues were then rinsed in phosphate buffered saline (PBS, pH 7.2), transferred into a cryoprotective solution (34.5% glycerol, 30% ethylene glycol, 20% PBS, 15.5% distilled water) and stored at –20≡C until further processing.

### Assessment of vestibular function in rats

To evaluate vestibular function, we assessed the tail-lift reflex as described (Martins-Lopes et al., 2019; Maroto et al., 2021 a,c). Briefly, rats are held by the base of the tail and quickly but gently lifted upwards to approximately 40 cm above ground and immediately returned down. In healthy rats, this triggers a trunk and limbs extension reflex in a landing response. Loss of vestibular function results in loss of the anti-gravity extension reflex, so the rat curls ventrally (Hunt et al., 1987; Pellis et al., 1991; Llorens et al., 1993; Maroto et al., 2021c). The reflex response of the animal is recorded from the side with high-speed video (we use 240 frames per second with a GoPro 5 camera) and the records are used to obtain the minimum angle formed by the nose, the back of the neck and the base of the tail during the tail-lift manoeuvre (tail-lift angle, TLA), as described (Martins-Lopes et al., 2019; Maroto et al., 2021a,c). The TLA has been demonstrated to provide an objective measure of permanent or reversible vestibular loss (Martins-Lopes et al., 2019; Maroto et al., 2021a,c).

### Electron microscopy

For SEM and TEM, rat vestibular epithelia were dissected in 2.5% glutaraldehyde in 0.1 M cacodylate buffer (pH 7.2), and fixed overnight in this solution. The epithelia were then rinsed with cacodylate buffer, post-fixed for 1 h in 1% osmium tetroxide in the same buffer, rinsed again and stored in 70% ethanol at 4≡C until further processing. Afterwards, the samples were dehydrated with increasing concentrations of ethanol, up to 100%. For TEM analysis, the lateral crista was embedded in Spurr resin. To select the level of interest, semi-thin sections (1 μm) were stained with 1% toluidine blue and examined in a light microscope. Ultrathin sections were stained with uranyl acetate and lead citrate and were observed with a JEOL 1010 microscope at 75-80 kV. For SEM analysis of epithelial surfaces, the dehydrated specimens were dried in a Polaron E3000 critical-point dryer using liquid CO2, coated with carbon, and observed in a JEOL JSM-7001F field emission SEM.

### Immunohistochemistry

The following commercial primary antibodies were used: mouse monoclonal anti-CASPR1 (clone K65/35, RRID: AB_2083496, used at 1:400 dilution) and anti-PSD95 (clone K28/43, RRID: AB_2292909, 1:100) from the UC Davis / NIH NeuroMab Facility; mouse monoclonal anti-Ribeye/CtBP2 (clone 16/CtBP2, BD Transduction Labs, RRID: AB_399431, 1:100); rabbit anti-myosin VIIa (25-6790, Proteus Biosciences, RRID: AB_10015251; 1:400); guinea pig anti-calretinin (214.104, Synaptic Systems, RRID: AB_10635160, 1/500); goat anti-SPP1 (AF808, RD systems, RRID: AB_2194992, 1/200). We also used a rabbit anti-KCNQ4 (PB-180, 1/1000) donated by Bechara Kachar (National Institute on Deafness and Other Communication Disorders, NIH; Beisel et al., 2005). To reveal the primary antibodies, we used Day-Light-405, Alex-488, Alexa-555 and Alexa-647-conjugated secondary antibodies from Invitrogen / Thermo Fisher. To image synaptic puncta, we used isotype-specific secondary antibodies, i.e., anti-mouse-IgG1 to reveal the Ribeye/CtBP2 antibody and anti-mouse IgG2a to reveal the PSD-95 antibody.

After dissection, the rat vestibular epithelia were fixed for 1 h at room temperature in 4 % paraformaldehyde in PBS. These samples were rinsed in PBS, transferred into the cryoprotective solution and stored at −20≡C. To circumvent potential bias from batch-to-batch differences in immunolabelling, we processed in parallel specimens from each experimental group. For immunolabelling, rat or human tissues were washed with PBS to remove the cryoprotective solution, permeated in 4%Triton-X-100 in PBS for 1 h at room temperature, and then blocked in 0.5% Triton-X-100 and 1% fish gelatine in PBS (Sigma-Aldrich, cat.# G7765) also for 1 h at room temperature. They were then incubated with the mixed primary antibodies in 0.1%Triton-X-100 and 1% fish gelatine in PBS for 24 h at 4≡C, followed by incubation with the secondary antibodies in the same conditions (Lysakowski et al., 2011). The samples were thoroughly rinsed with PBS following each incubation and gently rocked during the incubations and washes. The immunolabelled human epithelia were whole-mounted in Mowiol medium, whereas the rat epithelia were sectioned before mounting in the same medium. To obtain sections, the tissues were embedded in a mixture of 0.49% gelatin, 30% bovine serum albumin, and 20% sucrose in PBS overnight at 4 ≡C. This same medium was used to form a block by solidification with 2 % glutaraldehyde, one sample placed oriented in its surface, and a second block formed on top of it. Using a Leica VT1000S vibrating microtome, sections of 40 μm were obtained from this layered block with the specimen placed in between the two halves. A detailed version of this protocol has been published (Maroto et al., 2021b).

### Confocal microscopy observation and analysis

Tissue sections were observed in a Zeiss LSM880 spectral confocal microscope. In animal studies, image acquisition settings were maintained across samples from each immunolabelling batch. To obtain HC counts, Z-stacks of optical sections were obtained each 0.5 μm spanning 25 μm with a 63X objective. Then, we obtained 8 sequential images (3 μm thick each) by the maximum intensity projection of 6 consecutive images from the original stack. The ImageJ software (National Institute of Mental Health, Bethesda, Maryland, USA) was used for quantitative image analyses. Epithelia of animals injected once a day with streptomycin were labelled with anti-MYO7A, anti-CASPR1, and anti-calretinin antibodies. HCI were defined as MYO7A+ cells surrounded by a calyceal CASPR1+ label, even in cases where this label appeared thin and fragmented, or cells with an unambiguous HCI shape with a long and narrow neck. HCII were defined as cells labelled with both the anti-MYO7A and anti-calretinin antibodies. MYO7A+ cells not identified as HCI or HCII (3-6 %) were excluded from the analysis. For each animal and epithelial zone, a representative count of HCI and HCII was obtained by averaging counts from images 1, 4 and 7 from the series of 8 sequential images generated from a representative histological section. The histological sections used for counting were obtained from similar locations in all animals. To evaluate the loss of CASPR1 label in the calyces, the amount of fluorescence in the corresponding colour channel was measures in individual calyces using a 100 μm^2^ region of interest on a single optical section. For each animal, epithelium and zone, the average amount of fluorescence was obtained from all HCs available for analysis up to 40 cells. Data were normalized as percentage of the mean fluorescent intensity of control samples in the same batch of immunohistochemical processing and confocal imaging. Then, statistical analyses were performed to compare among groups of animals. To compare the distribution of KCNQ4 in the inner and outer membranes of the calyx, we assessed the profile of fluorescence intensity along a line drawn across the calyx as described (Sousa et al., 2009; Sedó-Cabezón et al., 2015). Peak fluorescence values were obtained for the inner and outer membrane of all calyces included in one representative section of the epithelium, and average values were obtained per animal. To count synaptic puncta, we imaged the tissues at 0.3 μm intervals to obtain Z-stacks of 2.7 μm. Only HC with their nuclei extending top to bottom of the stack were used for counting, so the counts obtained were from a thick slice from the middle region of the cell, not the whole cell. We counted presynaptic Ribeye/CtBP2 puncta, postsynaptic PSD95 puncta, and pairs of Ribeye and PSD95 puncta that overlapped or were closely apposed (co-localization). For each animal, average counts were obtained from a minimum of 30 HC.

Human samples were observed as whole mounts. The tissues were completely examined with 40X and 63X objectives. Representative Z-stacks, encompassing the entire thickness of the sensory epithelium, were obtained from both striola and peripheral regions. The full stacks were thoroughly studied to determine the presence of pathological features. This included colour channel splitting and merging, observation at several different contrast and brightness adjustments, 3D reconstructions, maximum intensity projection of stacks or sub-stacks and orthogonal projections. Images in Fig. 10 were prepared for illustration purposes only.

### Data analysis

Data show mean +/- standard error of the mean except where indicated. Body weight and behavioural data were analysed with repeated-measures MANOVA (Wilks’ criterion) with the day as the within-subject factor. Day-by-day analysis was performed after significant day-by treatment interactions were recorded. Other data were analysed by appropriate ANOVA designs and Duncan’s post-hoc test, or by Student’s t-test. The IBM SPSS Statistics 25 program package was used.

## RESULTS

### Effects of streptomycin sulphate on body weight and overall health

To study the sub-chronic toxicity of streptomycin on the vestibular system, we administered male and female rats with one daily dose of streptomycin sulphate (0, 100, 300, 400 or 500 mg/kg x day - groups CTRL, STR-100, STR-300, STR-400, STR-500) for up to 6 or 8 weeks, starting at day 21 of age. Other groups of rats were administered with two daily doses of 0, 300, 350 or 400 mg/kg - groups CTRL, STR-300×2, STR-350×2, STR-400×2). Rats administered with streptomycin showed an acute sedative effect starting within a few minutes after each daily dose, and was most evident in the group receiving the highest dose (STR-500): the animal stayed quiet with decreased responsiveness and with an overtly low muscle tone. This effect usually lasted 15-20 min but sometimes lasted for up to two hours after injection. During this sedation period, two rats died in the STR-500 group on days 5 and 36, likely from respiratory arrest. Additionally, two rats died for unknown reasons after days 20 and 37 in the STR-300 and STR-500 groups, respectively. No animals died when larger daily doses were administered via two daily injections. Recuperation from this sedative effect was complete and surviving animals showed normal behaviour except in regards to the evidences of vestibular dysfunction explained below. As shown in Suppl. Fig. S1, streptomycin animals showed an increase in body weight at a smaller rate than that shown by control rats. This resulted in significant differences in streptomycin versus control body weights after five or more weeks of one daily dose, and by two weeks in the groups receiving two daily doses.

### Loss and recuperation of vestibular function after sub-chronic streptomycin exposure

Sub-chronic exposure to streptomycin sulphate caused a dose-dependent loss of vestibular function, as indicated by the rat’s abnormalities in spontaneous motor behaviour. Vestibular deficient animals showed ataxia, spontaneous circling, and head bobbing. The loss of function was assessed using high-speed video records of the tail-lift reflex as described (Martins-Lopes et al., 2019; Maroto et al., 2021a,c). A similar dose-dependent effect occurred in male and female animals (Suppl. Fig. S2), and therefore animals of both sexes were pooled for all further analyses. The decrease in tail-lift angles, indicative of loss of vestibular function, was not significant in the STR-100 group, and smaller in the STR-300 and STR-400 groups compared to the STR-500 group (Suppl. Fig. S3), so further analysis on the one daily dose paradigm focused on the 500 mg/kg x day dose level. As shown in Figure 1A, a progressive reduction in tail-lift angles was recorded in the STR-500 rats. Repeated-measures MANOVA of the tail-lift data from days 0 (pre-test) to 42 resulted in significant day (F[6,28]=15.38, p<0.001), treatment (F[1,33]=18.0, p<0.001), and day X treatment interaction (F[6,28]=9.766, p<0.001) effects, with significant differences between CTRL and STR-500 mean values recorded at 4 to 6 weeks of exposure. Extension of the administration up to 8 weeks caused only a modest additional effect on vestibular function (Suppl. Fig. S4A).

**Figure 1.**
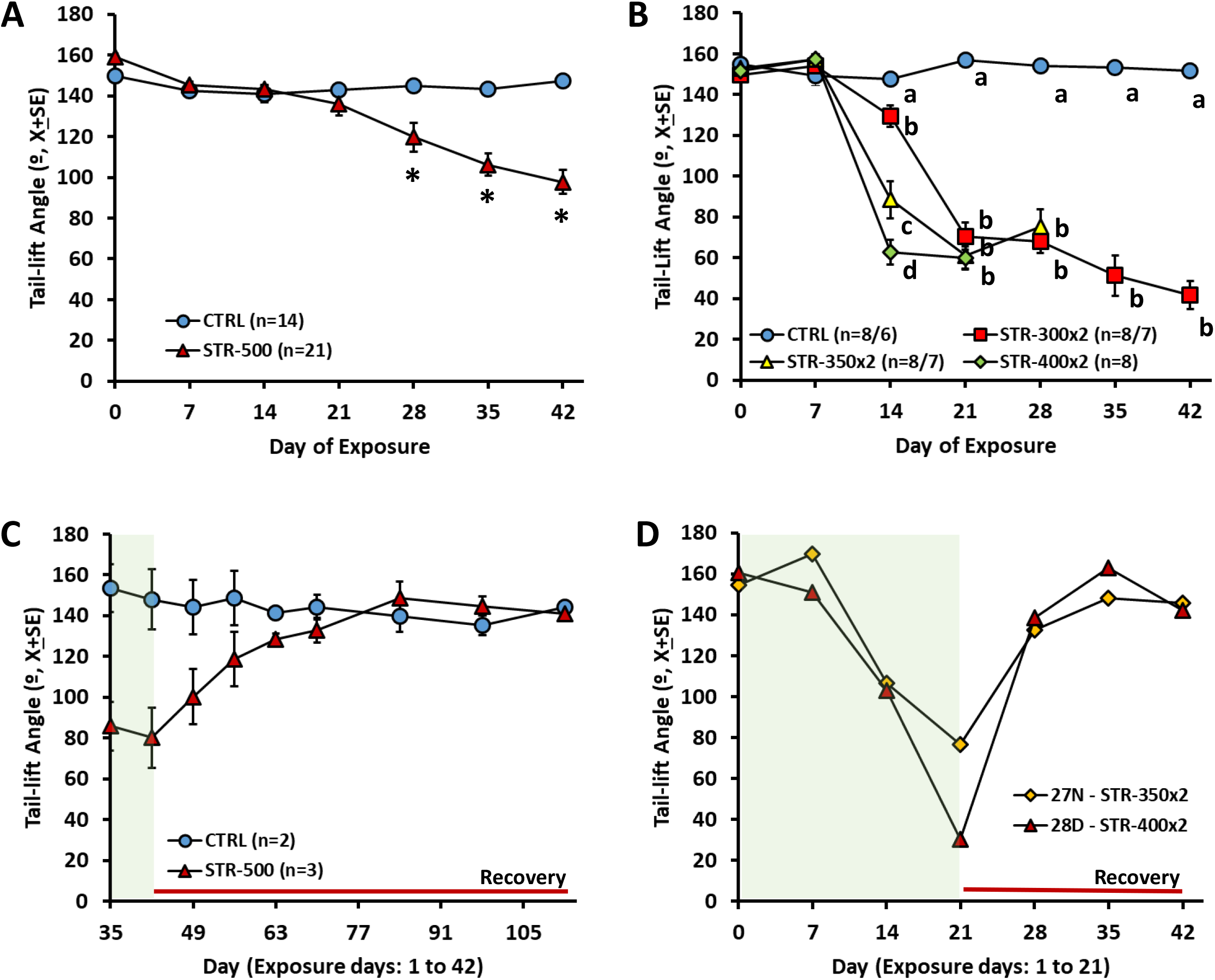
Effects of sub-chronic streptomycin on the tail-lift reflex of rats. Data are mean + SE minimum angles formed by the nose, the back of the neck, and the base of the tail during the reflex behaviour elicited by a tail-lift manoeuvre. Loss of vestibular function causes loss of the anti-gravity trunk extension reflex, which results in ventral curling and thus in smaller angles. A. Effects of one injection per day at 0 (CTRL, vehicle control) or 500 mg/kg (STR-500) of streptomycin for 6 weeks, starting at 21 days of age. * p<0.05, different from control group after significant day by treatment interaction effect in repeated-measures MANOVA. B. Effects of two injections per day at 0 (CTRL, vehicle control), 300 (STR-300×2), 350 (STR-350×2), or 400 (STR-400×2) mg/kg of streptomycin for 6, 6, 4, or 3 weeks, respectively, starting at 21 days of age. The numbers of animals were reduced to 6 in the CTRL group after day 28, and to 7 in the STR-300×2 and STR-350×2 groups after day 21. **a, b, c, d:** groups with different letters are significantly different (p<0.05), Duncan’s test after significant ANOVA on that day. C. Recuperation of the tail-lift reflex after the end of exposure in rats administered once a day with 500 mg/kg (STR-500) of streptomycin for 6 weeks. D. Loss and recuperation of the tail-lift reflex in two individual rats administered twice a day with 350 (STR-350) or 400 mg/kg (STR-400) of streptomycin for 3 weeks, followed by 3 weeks of recovery.

By administering streptomycin in two daily injections, larger doses could be given to the animals and circumvent the risk of death associated with the acute sedative effects of the drug. As shown in Fig. 1B, all the STR-300×2, STR-350×2, and STR-400×2 groups showed faster and deeper loss of vestibular function than the STR-500 group (Fig. 1A). MANOVA of the tail-lift data from the two daily injections experiment up to day 21 resulted in significant day (F[3,26]=191.1, p<0.001), treatment (F[3,28]=45.89, p<0.001), and day X treatment interaction (F[9,63.4]=20.92, p<0.001) effects. The effect was dose-dependent, and larger doses caused a faster progression of the decline in vestibular function.

To evaluate the potential for functional recuperation, some animals were allowed to survive after the end of the sub-chronic streptomycin exposure. As shown in Fig. 1C, rats exposed to 500 mg/kg · day of streptomycin for 6 weeks regained normal tail-lift angles during the recovery period. Functional recuperation was also observed in the STR-500 rats exposed for 8 weeks (Suppl. Fig. S4B). The individual values of animals allowed to recover after 3 weeks of two daily doses of streptomycin also showed a clear recuperation of the tail-lift reflex (Fig. 1D).

### Sub-chronic streptomycin exposure caused partial loss of HCs in the vestibular sensory epithelia

To evaluate the major effects of streptomycin on the vestibule, we observed surface preparations of the sensory epithelia using scanning electron microscopy (SEM). As reported in previous studies (Llorens et al., 1993; Seoane et al., 2001a; Sedó-Cabezón et al., 2015), SEM observation of control epithelia (n=16) (Fig. 2A; Suppl. Fig. S5A, B) revealed morphological features matching literature descriptions of healthy untreated rats with high densities of stereocilia bundles. Epithelia from STR-300 animals (6 weeks of exposure, n=3) showed similar features to control epithelia, except for indications of a few missing hair bundles or occasional bundles with fused stereocilia (data not shown). For animals administered with larger doses of streptomycin once a day, the SEM observations revealed scant to moderate loss of stereocilia bundles. Similar findings were obtained in STR-500 rats with small difference observed between animals exposed for 6 (n=6) or 8 (n=7) weeks, and in STR-400 rats exposed for 8 weeks (n=2). Hair bundle loss was roughly similar in cristae (Suppl. Fig. S5C), utricles (Suppl. Fig. S5D), and saccules (Fig 2B, C), with no evidence of an overt difference in susceptibility among receptors. In all cases, the loss of stereocilia bundles was partial and many bundles were still present in the sensory receptors of the worst cases (Suppl. Fig. S5E,F). In these SEM observations, images indicating ongoing damage were scarce. Nevertheless, some evidence for fused stereocilia and HC protrusion were found (Fig 2D,E). Animals given two daily injections of streptomycin displayed a larger loss of stereocilia bundles than the animals administered for once a day (Suppl. Fig. S6). Evidence for ongoing damage was scarce in some epithelia, but clear in other samples, including images of epithelial scars, bundle fusion, and cell protrusion towards the endolymphatic cavity (Suppl. Fig. S6). The average extent of hair bundle loss was similar in STR-300×2 rats treated for 6 weeks (n=7), STR-350×2 rats treated for 4 weeks (n=6), and STR-400×2 rats treated for 3 weeks (n=6).

**Figure 2.**
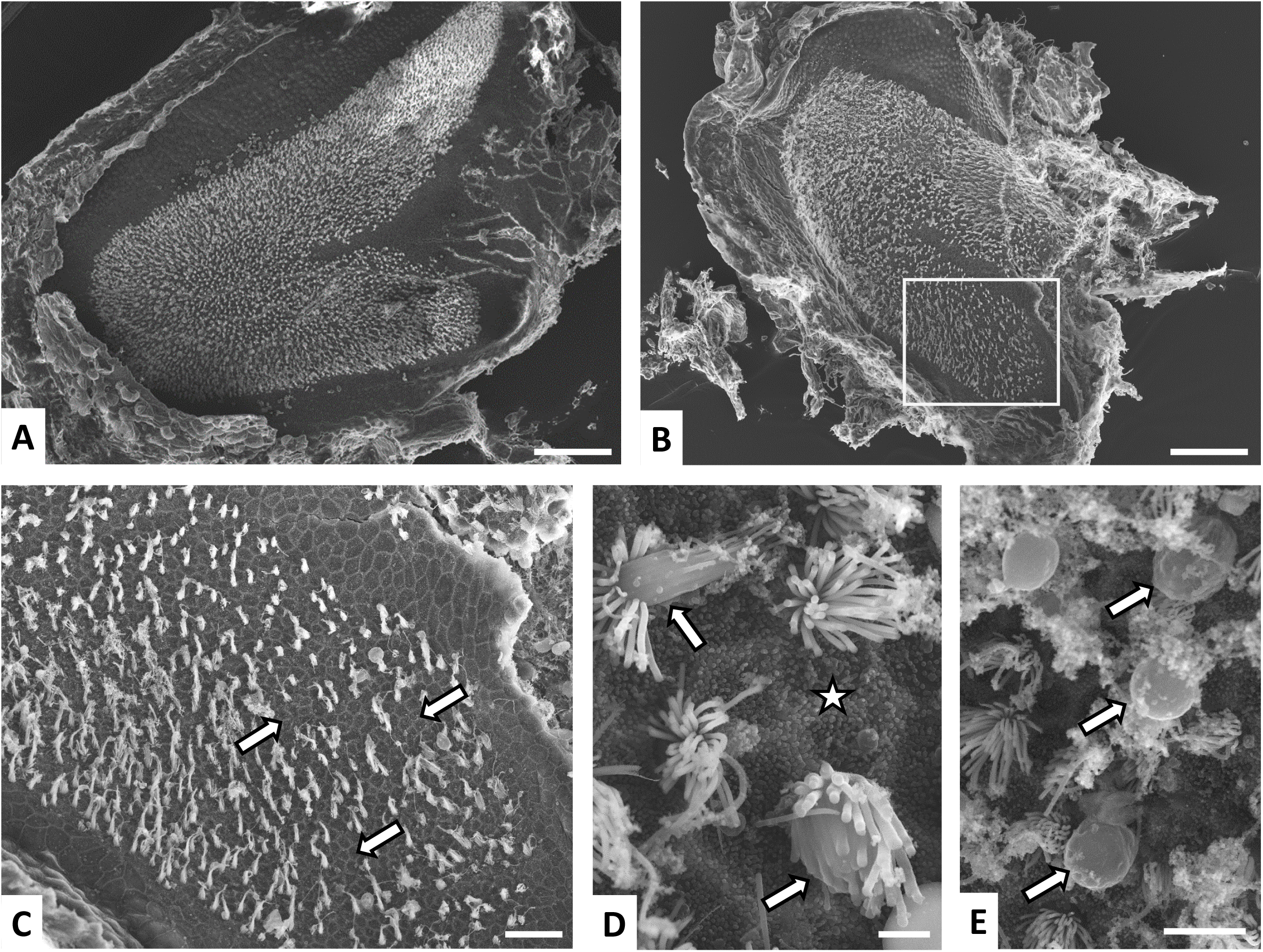
Effects of sub-chronic streptomycin on the vestibular sensory epithelium of the rat as observed in surface preparations of saccules examined by SEM. A. Control saccule, showing a high and uniform density of stereocilia bundles throughout the macula. B. Effect of one injection per day of 500 mg/kg of streptomycin (STR-500) for 8 weeks. A reduced density of stereocilia bundles is noticeable at this low magnification. The boxed area is shown at higher magnification in panel C. C. Higher magnification of the saccule shown in panel B. Note the areas where the lack of stereocilia bundles is evident (arrows). **D and E.** High magnification views of a saccule from a STR-500 rat at 6 weeks of treatment. Note the contacts between the apical edges of supporting cells, marked by a high density of microvellosities, denoting the lack of stereocilia bundles (star), and the bundles showing stereocilia fusion and HC protrusion into the luminal cavity (arrows). **Scale Bars:** 100 μm in **A** and **B;** 20 μm in **C**; 2 μm in **D**; 5 μm in **E**.

To determine the loss of HCs revealed by the decreased densities of hair bundles observed by SEM in STR-500 rats, we obtained type I HC (HCI) and type II HC (HCII) counts from the vestibular epithelia of the other ear of the same animals. To this end, we immunostained the epithelia with antibodies against myosin-7a (MYO7A), CASPR1, and the calcium-binding protein calretinin, revealed by fluorescent secondary antibodies. In the confocal microscopy images, HCI were identified as MYO7A+ cells (a pan-HC marker, Hasson et al., 1997) surrounded by a CASPR1+ calyx (Sousa et al., 2009; Lysakowski et al., 2011; Sedó-Cabezón et al., 2015), even if weak or fragmented, or with an unambiguous HCI shape with a long and narrow neck. As an estimate of HCII, we counted cells with double MYO7A and calretinin label (Dechesne et al., 1991). The data obtained, presented in Fig. 3A-C, reveal a significant decrease in HCI counts in the central region of the crista, in both the centre and periphery of the utricle and in the periphery of the saccule. ANOVA results were: central crista, F[3,27]= 11.95, p<0.001; peripheral crista, F[3,24]=2.76, p=0.064; central utricle, F[3,25]=17.36, p<0.001; peripheral utricle, F[3,26]=23.51, p<0.001; central saccule, F[3,20]=1.68, p=0.20-N.S.; peripheral saccule, F[3,22]=9.08, p=0.001. The most affected receptor was the utricle with a 52-67 % decrease in HCI counts in STR-500 animals, followed by the crista, where a 30-40 % loss was recorded. In both the crista and the utricle, similar effects were recorded in animals exposed for 6 weeks and animals exposed for 8 weeks. In the saccule, the loss was smaller than in the crista and utricle receptors, and greater at 8 weeks than at 6 weeks of treatment. HCI counts remained reduced in animals allowed to recover after the end of the exposure period, indicating an insignificant capacity for regeneration. In contrast to HCI, no significant differences were obtained among treatment groups in HCII counts (all p values > 0.1).

**Figure 3.**
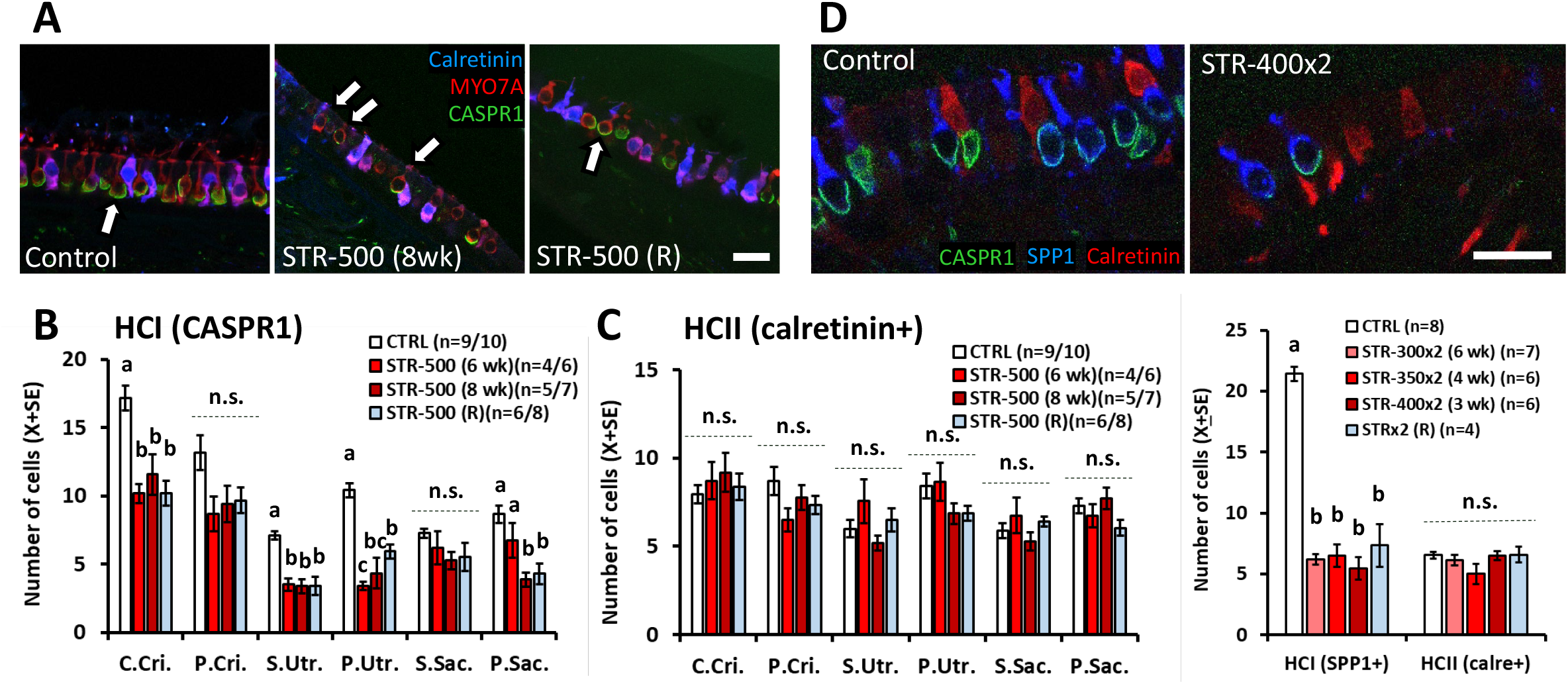
Effect of sub-chronic streptomycin on the numbers of HCI and HCII in the vestibular epithelia of the rat. **A, B, C.** Effect of 500 mg/kg of streptomycin, once a day, for 6 or 8 weeks, as recorded at the end of the exposure period or at the end of a recovery period of 8 weeks after treatment. Vestibular cristae, utricles and saccules were immunolabelled with anti-MYO7A (red, to label all HCs), anti-CASPR1 (green, to identify HCI) and anti-calretinin (blue, to label HCII) antibodies, and examined by confocal microscopy. The central/striola and peripheral regions of the epithelia were evaluated separately. A. Representative images from the peripheral utricle of a vehicle control rat, a rat treated with streptomycin for 8 weeks (STR-500 8wk), and a recovery (STR-500 R) rat. HCI were defined by MYO7A+ cells with adjacent CASPR1 label, even if thin or fragmented, and no calterinin label (arrows). Note the fragmentation of the CASPR1 label in the STR-500 (8wk) rat. **Scale bar:** 20 μm. **B and C.** Counts of HCI **(B)** and HCII **(C)** cells. Data are mean number of cells +/- SE per section. For each animal and epithelial zone, numbers were obtained by averaging counts from 8 confocal stacks from a representative histological section. **D, E.** Effect of 300 (STR-300×2), 350 (STR-350×2) or 400 (STR-400×2) mg/kg of streptomycin, twice a day, for 6, 4, or 3 weeks. The recovery (R) group includes animals allowed 3 weeks of recovery after receiving STR-350×2 or STR-400×2 for 3 or 4 weeks. Vestibular cristae were labelled with anti-SPPl (blue, to label HCI), anti-CASPR1 (green) and anti-calretinin (red) antibodies. HCI were defined as SPP1+ cells and HCII were cells with calretinin label. **D.** Partial view of representative sections of control and streptomycin cristae. **Scale bar:** 20 μm. **E.** Relative numbers of HCI and HCII in the crista sections. Data are mean number of cells +/- SE per transversal section, including central and peripheral regions. In **B, C,** and **E,** the number of replicas in the figure legends indicate minimum/maximum number of animals counted per epithelium and zone in the treatment group, **a, b, c, ab:** groups not sharing a letter are different (p<0.05), Duncan’s multiple-comparisons test after significant (p<0.05) one-way ANOVA. **(n.s.):** non-significant (p>0.05) group differences in the ANOVA analysis.

As explained above, the SEM observations indicated a loss of HC after two daily injections of streptomycin greater than that recorded after one daily injection. To estimate HC loss according to cell type, we immunostained crista receptors with antibodies against secreted phosphoprotein 1 (SPP1) / osteopontin, CASPR1, and calretinin. HCI were identified by the SPP1 label (Burns et al., 2015; McInturff et al., 2018; Maroto et al., 2021b), while the main population of HCII was identified by the calretinin+ label. As shown in Fig. 3D,E, the number of HCI was greatly reduced after two daily injections of streptomycin (F[4,26]=77.64; p<0.001), while the number of HCII was unchanged (F[4,26]=1.48, p=0.24). The loss of HCI was similar in the three groups of rats treated with streptomycin (STR-300×2, STR-350×2, STR-400×2), and no recuperation in HCI counts was recorded in the recovery group.

### Calyceal junction dismantlement, afferent fragmentation, and synaptic loss after sub-chronic streptomycin exposure

In addition to HCI loss, rats exposed to sub-chronic streptomycin displayed alterations in the remaining HCIs and their afferents. In rats exposed to streptomycin once a day (STR-500), a reversible loss of CASPR1 fluorescence occurred in the calyces contacting the surviving HCIs (Fig.5). To assess this effect, we measured the amount of CASPR1 fluorescence associated with the remaining individual calyces. The loss and recuperation of the CASPR1 label was found across the three epithelia and in both central/striolar and peripheral zones. All ANOVA comparisons resulted in significant differences (all F[3, 19 to 27] > 5.3, all p’s<0.006). As shown in Fig. 4, the confocal microscope images also revealed a reversible decrease in MYO7A fluorescence intensity in the HCs of the STR-500 animals. This effect was quantified in a subset of samples and significant group differences were found (F[2,11]=5.0, p=0.029). The post-hoc analysis confirmed that the label (in arbitrary units) was reduced from 9583 + 1290 (X+SE, n=5) in CTRL animals to 6232 + 781 (n=4) in STR-500 (8 wk) animals (p<0.05, Duncan’s test), and regained normal level (10522 + 621, n=5) after recovery. In animals given two injections a day, the intensity of CASPR1 immunoreactivity in the calyces contacting surviving HCI was also reduced by streptomycin (Suppl. Fig. S7).

**Figure 4.**
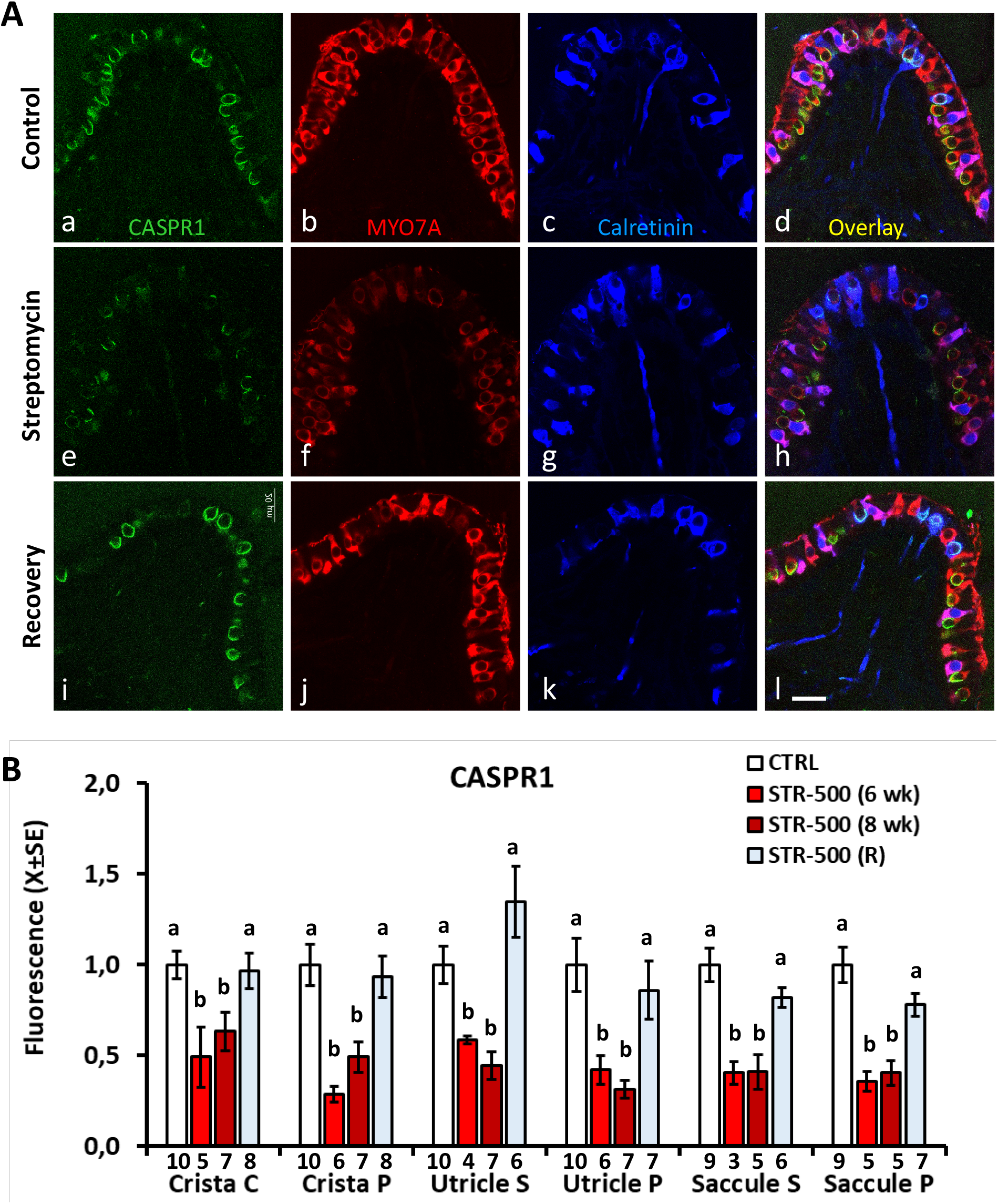
Effects of sub-chronic streptomycin on the expression of CASPR1 in the central/striola (C/S) and peripheral (P) regions of the vestibular crista, utricle and saccule of the rat. The samples were labelled with anti-MYO7A (red), anti-CASPR1 (green) and anti-calretinin (blue) antibodies. **A.** Representative images from mid-crista sections from a CTRL rat **(a** to **d),** a rat treated with 500 mg/kg of streptomycin, once a day, for 6 weeks **(e** to **h),** and a rat allowed a recovery period of 10 weeks after the streptomycin treatment **(i** to I). Note the loss and fragmentation of the CASPR1 label caused by streptomycin and its recuperation after recovery. The intensity of MYO7A label also decreased and recovered. B. Quantitative analysis of the CASPR1 fluorescence. Data are normalised arbitrary fluorescence amount per cell +/- SE. For each animal, epithelium and zone, the average amount of fluorescence was obtained from all HCs available for analysis up to 40 cells. Numbers below the bars indicate number of animals included per group, **a, b:** groups not sharing a letter are different (p<0.05), Duncan’s multiple-comparisons test after significant (p<0.05) one-way ANOVA.

**Figure 5.**
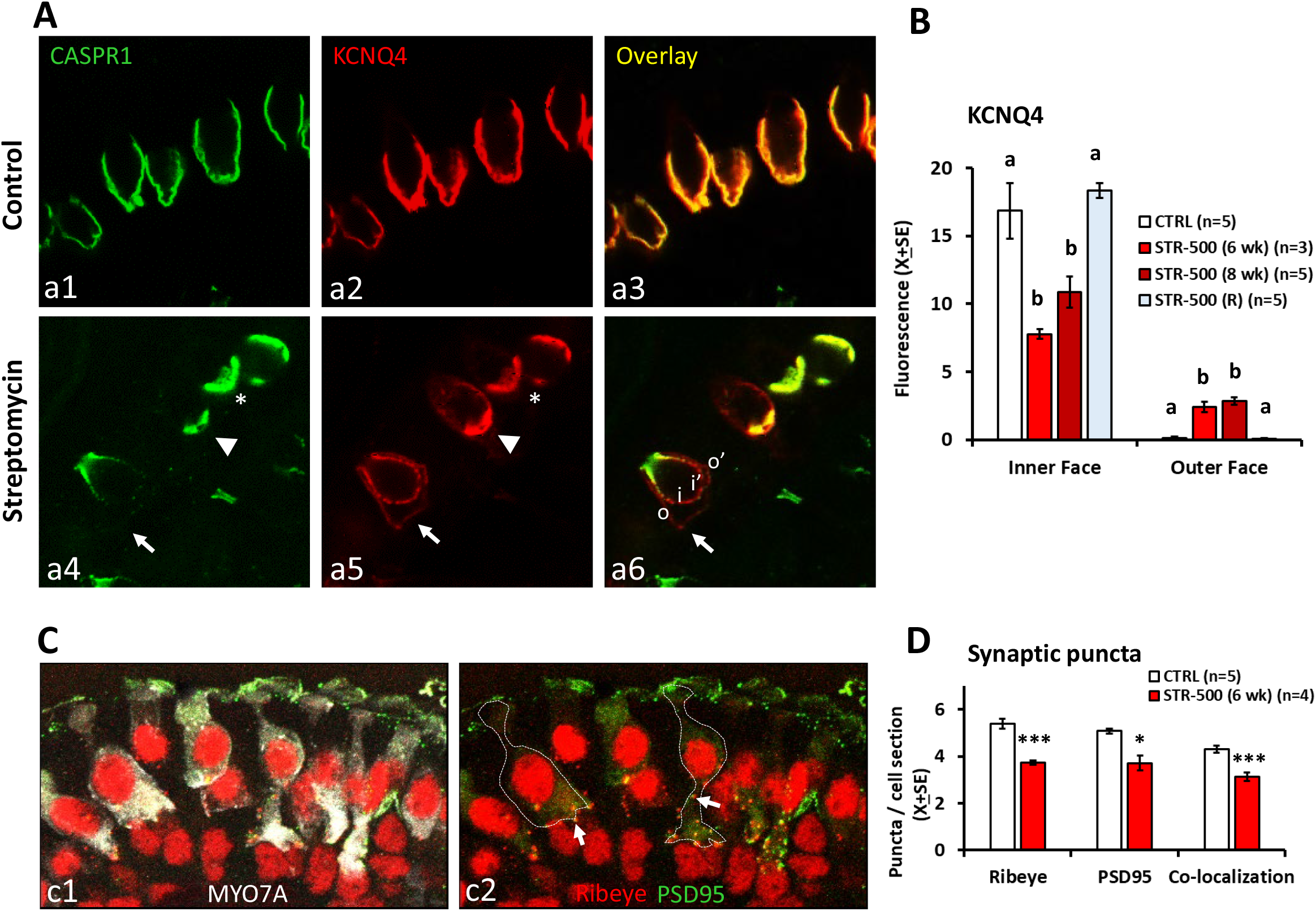
Effects of sub-chronic streptomycin on the expression of synaptic proteins in the HCI and calyx afferents of the rat. A. Redistribution of KCNQ4 in the inner and outer membranes of the calyx terminals afferent to HCI. Vestibular crista from control (a1-a3) and streptomycin (a4-a6) animals are shown after labelling with anti-CASPR1 (green) and anti-KCNQ4 (red) antibodies. Note the precise colocalization of CASPR1 and KCNQ4 in the control epithelium. After streptomycin, KCNQ4 mislocalisation concurs with CASPR1 loss. The asterisk indicates a calyx with a control-like co-localisation of CASPR1 and KCNQ4 in the inner membrane of the calyx. The arrowhead points to a calyx with CASPR1 loss and reduced co-localisation, with the KCNQ4 antibody labelling a greater area than the CASPR1 antibody. The arrow points to the outer membrane of a calyx displaying greatly reduced CASPR1 label in the inner membrane and redistribution of KCNQ4 throughout both the inner (i and i’) and outer (o and o’) membranes. B. Quantitative analysis of the distribution of KCNQ4 label in the inner and outer face of the calyces in crista from control rats (CTRL), rats treated with 500 mg/kg of streptomycin (STR-500), once a day, for 6 or 8 weeks, and rats allowed a recovery period of 8-10 weeks after the streptomycin treatment. Note that control and recovery animals do not display KCNQ4 in the outer face membrane. Data are mean arbitrary fluorescence units +/- SE. The number of replicas indicate the number of animals included in the analysis per group, **a, b:** groups not sharing a letter are different (p<0.05), Duncan’s multiple-comparisons test after significant (p<0.05) one-way ANOVA. C. Labelling of pre-synaptic (Ribeye, red) and post-synaptic (PSD-95, green) proteins on HCs (MYO7A, white) in vestibular crista. In subpanel **c1,** the MYO7A label is included to show HC bodies in a control crista. Subpanel **c2** shows the same image without the MYO7A label but including lines delineating the limits of two HCs and arrows pointing to examples of pairs of ribeye and PSD-95 puncta. **D.** Effects of sub-chronic streptomycin on the numbers of synaptic puncta per 2.7 μm-tick optical sections of HCs. Data are mean number of pre-synaptic and post-synaptic puncta, as well as pre- and post-synaptic pairs (Co-localization), +/- SE, from control rats (CTRL) and rats treated with 500 mg/kg of streptomycin (STR-500), once a day, for 6 weeks. The number of replicas indicate the number of animals included in the analysis per group. *: p< 0.05, and ***: p<0.001, CTRL versus STR-500, Student’s T-test.

We also evaluated whether sub-chronic streptomycin altered the distribution of the potassium channel, KCNQ4, in the calyx membrane, as identified in CASP1-KO mice (Sousa et al., 2009) and in rats exposed to sub-chronic IDPN (Sedó-Cabezón et al., 2015). The data, shown in Fig. 5A,B, indicated that CASPR1 loss is associated with KCNQ4 redistribution in the calyces of the STR-500 rats. In control rats, the KCNQ4 label is found almost exclusively in the inner membrane, that is, in the calyceal junction area, and is absent in the outer membrane of the calyx. After streptomycin treatment, the immunofluorescence was reduced in the inner membrane, whereas significant labelling appeared in the outer membrane of the calyx. Therefore, KCNQ4 was abnormally distributed after streptomycin. ANOVA analyses of the amount of fluorescence in the inner and outer membranes were significant (F[3, 14]=12.1, P<0.001, and F[3,14]=49.9, p<0.001, respectively). Post-hoc analyses confirmed that streptomycin caused KCNQ4 redistribution, and that this effect was no longer present in the recovery animals.

Sub-chronic streptomycin also caused a reduction in the number of synaptic puncta as revealed by labelling with antibodies against ribeye/CtBP2, the core protein of the presynaptic ribbon, and against the post-synaptic scaffold protein PSD-95 (post-synaptic density 95). The mean number of puncta counts per cell in 2.7 μm thick sections of HCs was significantly reduced in STR-500 animals exposed for 6 weeks compared to CTRL animals (Fig. 5C,D). T-test comparisons resulted in p <0.001, p=0.018, and p=0.001 for ribeye, PSD-95, and co-localization (as defined in the methods section) values, respectively.

The effect of sub-chronic streptomycin on the calyceal junction was also studied at the ultrastructural level by TEM observation of samples from control rats (4 lateral crista, 2 saccules, 1 utricle), STR-500 rats exposed for 6 (5 lateral crista) or 8 (4 lateral crista) weeks, and STR-300×2 rats exposed for 6 weeks (3 saccules and 1 utricle). In control specimens, calyceal junctions were prominent with virtually no disruptions in the two basal thirds of contact between HCI and its afferent calyx (Fig. 6A,C). This junction consists of highly electron-dense pre- and post-synaptic membranes, a highly regular inter-membrane distance, and an electron-dense extracellular space. In contrast, many calyces in the streptomycin rats showed from fragmented to only partial to absent junctions (Fig. 6B, D-F). In these units, the membrane density was similar to that of regular membranes and the area displayed a more-variable inter-membrane distance, as well as a reduced extracellular electron density. In addition, fragmented calyces were also identified in the samples from the treated animals, and these showed absence of the calyceal junction (Fig. 6E, F). No fragmented calyces were observed in the control epithelia.

**Figure 6.**
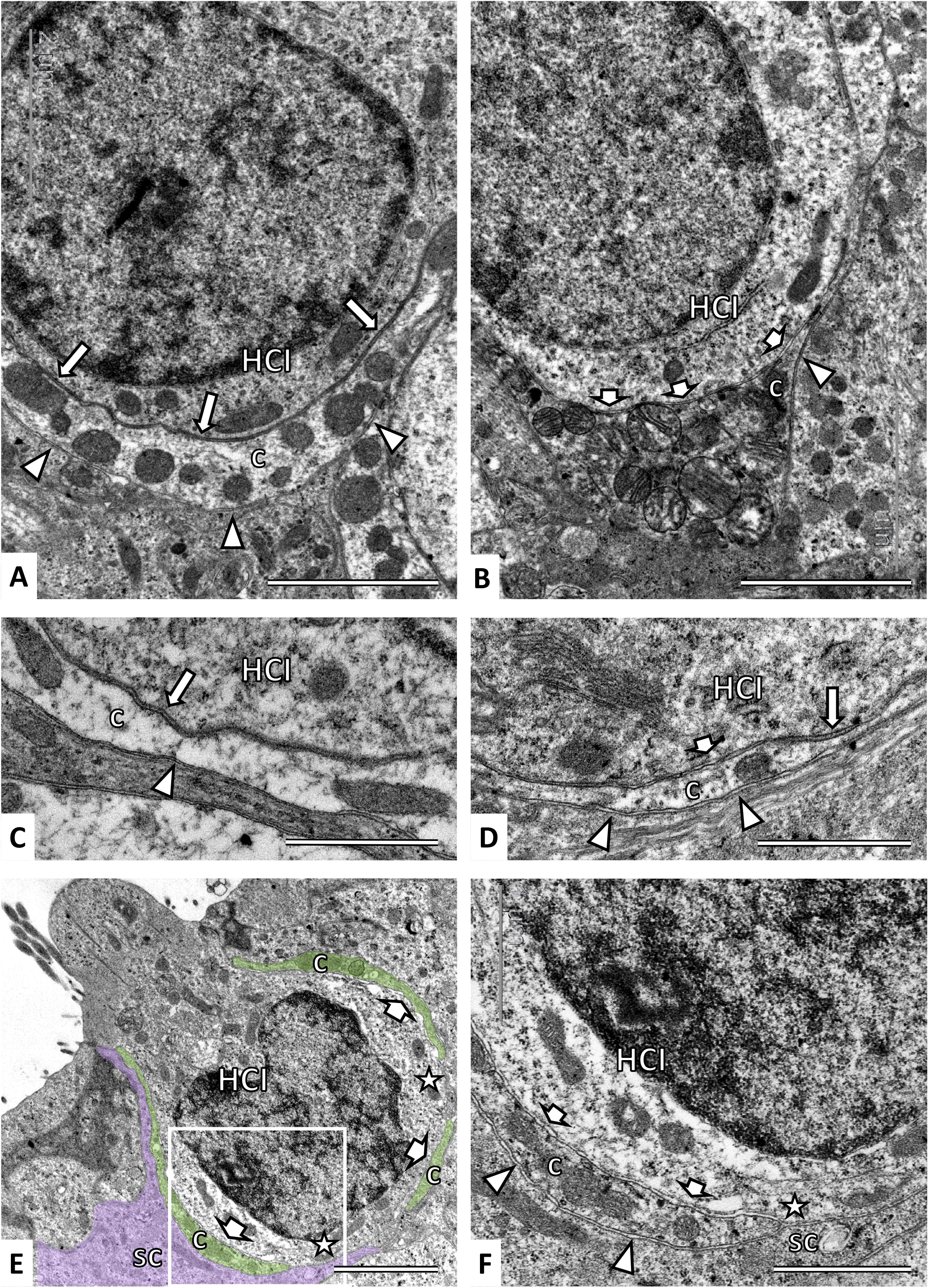
Effects of sub-chronic streptomycin on the calyceal junction and calyx integrity as observed by TEM. **A.** Control crista. The calyceal junction is revealed by the high electron density of the apposing membranes (long arrows) of HCI and inner side of the calyx (c), higher than that found in the outer side of the calyx (arrowheads). **B.** Crista of a rat exposed to 500 mg/kg of streptomycin, once a day, for 6 weeks. In most of the area where the calyceal junction should be found, the apposing membranes show a low electron density (short arrows), similar to that in the outer face of the calyx (arrowhead). C. Higher magnification of the calyceal junction (long arrow) and the outer side of the calyx (arrowhead) in a control saccule. D. Higher magnification from a crista of a rat exposed to 500 mg/kg of streptomycin, once a day, for 6 weeks. Note the short segment of calyceal junction (long arrow) that remains together with areas devoid of it (short arrow) and the similarity between the latter and the membranes in the outer side of the calyx (arrowheads). E. Calyceal junction dismantlement and calyx fragmentation in a HCI showing evidence of ongoing extrusion, from a utricle of a rat exposed to 300 mg/kg of streptomycin, twice a day, for 6 weeks. Note the absence of calyceal junction (short arrows), the fragmentation of the calyx (c, green shadow), and the direct contact between supporting cells and the HCI (stars). The purple shadow highlights one supporting cell (sc). F. Higher magnification of the boxed area in panel E. Note the similarity of the membranes between the place where the calyceal junction is missing (short arrows) and the outer side of the calyx (arrowheads). Note also the direct contact between a supporting cell (sc) and the HCI (star). **Scale Bars: A, B,** and **E,** 2 μm; **C, D,** and **F,** 1 μm.

### Hair cell extrusion after sub-chronic streptomycin exposure

The occurrence of HC extrusion was evaluated by immunofluorescent analysis in samples from animals exposed to streptomycin twice a day. The MYO7A and CASPR1 antibodies were combined with an anti-radixin antibody to label the apical microvilli of the supporting cells and thus outline the surface of the epithelium. Images indicating HC extrusion were not obtained in most sections of the crista (n=4 CTRL, 7 STR-300×2, 4 STR-350×2, and 4 STR-400×2) and saccule (n=2 CTRL, 7 STR-300×2, 5 STR-350×2, and 6 STR-400×2). However, as illustrated in Fig. 7A, protruding/extruding HCs were a common finding in most utricle sections from animals exposed to STR-300×2 for 6 weeks (n=7) or STR-350×2 for 4 weeks (n=5), although not in utricles from animals receiving STR-400×2 for 3 weeks (n=6). Control utricles (n=2) also showed no HC extrusion. In the STR-300×2 and STR-350×2 utricles, the occurrence was around 1-3 protruding/extruding HC per section (130-190 μm long, 25 μm thick).

**Figure 7.**
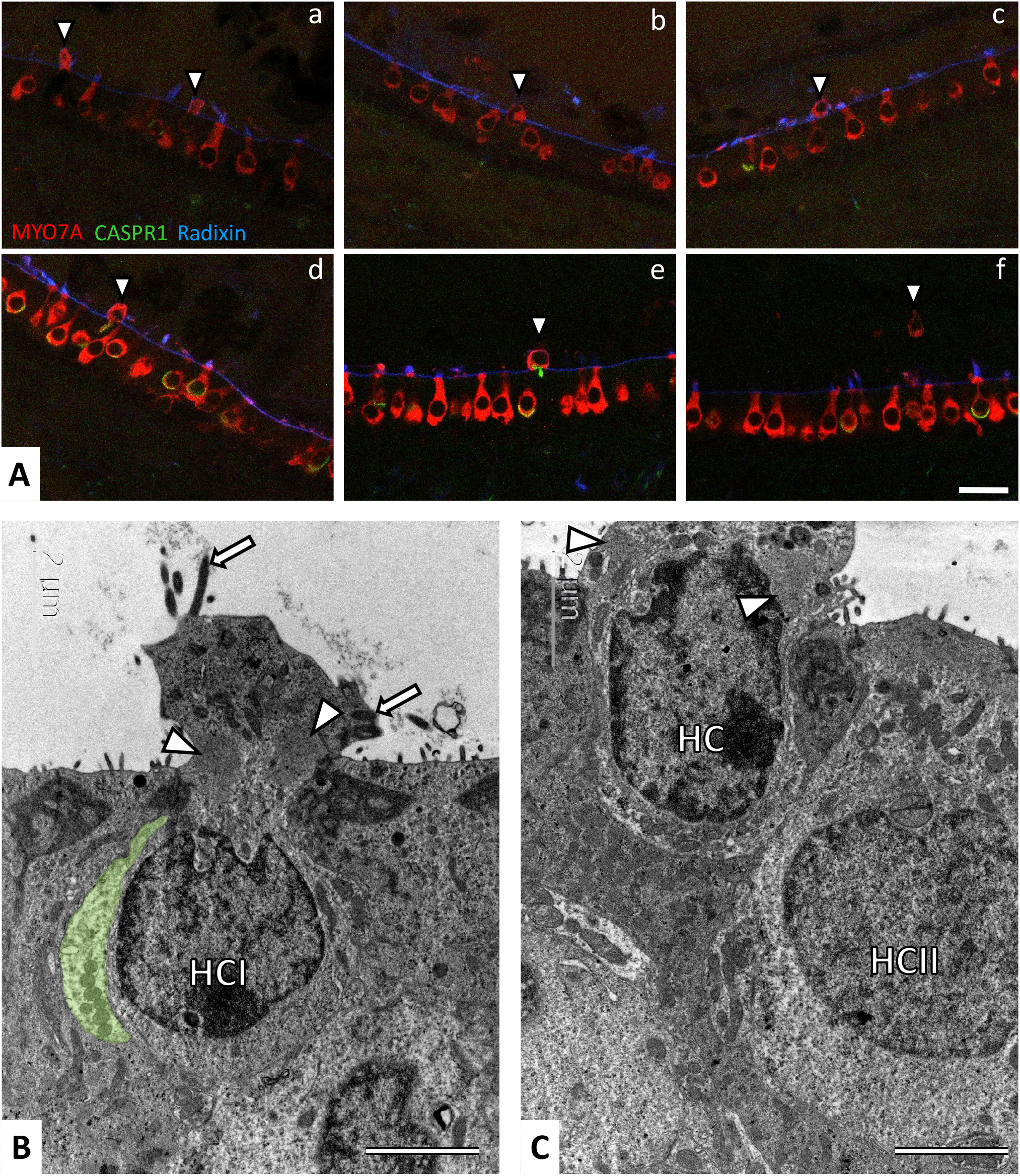
Evidence of HC extrusion in utricles of rats exposed to sub-chronic streptomycin as observed by immunofluorescence and TEM. The images are from rats administered twice a day with 350 mg/kg of streptomycin for 4 weeks, or with 300 mg/kg for 6 weeks. A. Utricles immunolabelled with antibodies against MYO7A (red), CASPR1 (green) and radixin (blue). The radixin label of stereocilia and microvilli outlines the surface of the epithelium. Arrowheads point to HCs protruding **(a, b),** extruding **(c, d, e)** or completely extruded **(f)** into the luminal cavity of the utricle. **B** and **C.** TEM images of HCs in a protrusion stage corresponding to that shown in panels **a** and **b,** respectively, of part **A.** Stereocilia (arrows) and striated organelles (arrowheads) identify the extruding cells as HCs. In B, note the position of the nucleus of the HC inside the epithelium, and the remnant of a calyx terminal (green shadow) that identifies the HC as a HCI. In **C,** note the position of the nucleus of the HC (unidentified type), half way into the luminal cavity of the utricle. **Scale Bars: A,** 20 μm; **B,** and **C,** 2 μm.

To corroborate that the extruding MYO7A+ profiles found in the immunofluorescent analyses corresponded to living HCs, we examined two utricles by TEM. As shown in Fig. 7B,C, the extruding HC showed displacement of the whole cell towards the lumen of the utricle. The ultrastructural features of the cytoplasm, organelles, and nucleus indicated continued activity of the cell.

### Occurrence of calyceal junction dismantlement in human vestibular epithelia

To evaluate whether calyceal junction dismantlement also occurs in human vestibular epithelia under stress, we studied samples from patients submitted to therapeutic labyrinthectomy or translabyrinthine tumour excision. The vestibular epithelia were immunolabeled with anti-CASPR1 antibodies combined with a variety of other antibodies, and a total of 29 samples yielded useful information. MYO7A, calretinin, and CASPR1 labels matched the labelling features described above in rat samples. Also, when tested together (n=9), CASPR1 and the extracellular matrix protein tenascin-C were found to localise together in the calyceal junction area, as described previously in rodent samples (Lysakowski et al., 2011; Sedó-Cabezón et al., 2015; Greguske et al., 2019). In contrast, the observed label of HCI by SPP1 was unreliable in the tested conditions. Among the 29 epithelia, HC densities varied from high, similar to that observed in samples from control rats, to almost null. Reduced HC density could result from both the patient’s age and vestibular pathology; the evaluation of this pathology was beyond the scope of the present study. Regarding the calyceal junctions, 18 samples were evaluated to have intact or mostly intact CASPR1-labelled calyces. These included 2/2 cases of Ménière’s Disease (Fig. 8A), 4/4 cases of meningioma (Fig. 8B,C), 1/1 case of an endolymphatic sac tumour, and 11/22 cases of vestibular schwannoma. Two other samples displayed very few HCs with only one CASPR1+ or none, hampering the achievement of any conclusion on calyceal junctions. However, 9 of the samples, all of them from vestibular schwannoma patients, provided images suggesting that an active process of calyceal junction dismantlement was taking place. This included calyces with patched CASPR1 labelling (Fig 8D-F) or lack of CASPR1 in calyx-only units (Fig. 8G). The patched feature of the label in some calyces was confirmed in 3D reconstructions (Suppl. Fig. S8). The number of days the samples were in fixative does not explain this effect, as it was observed in tissues fixed for 1 day (n=5), 4 days (n=1), 7 days (n=1), and unrecorded (but less than 8) days (n=2).

**Figure 8.**
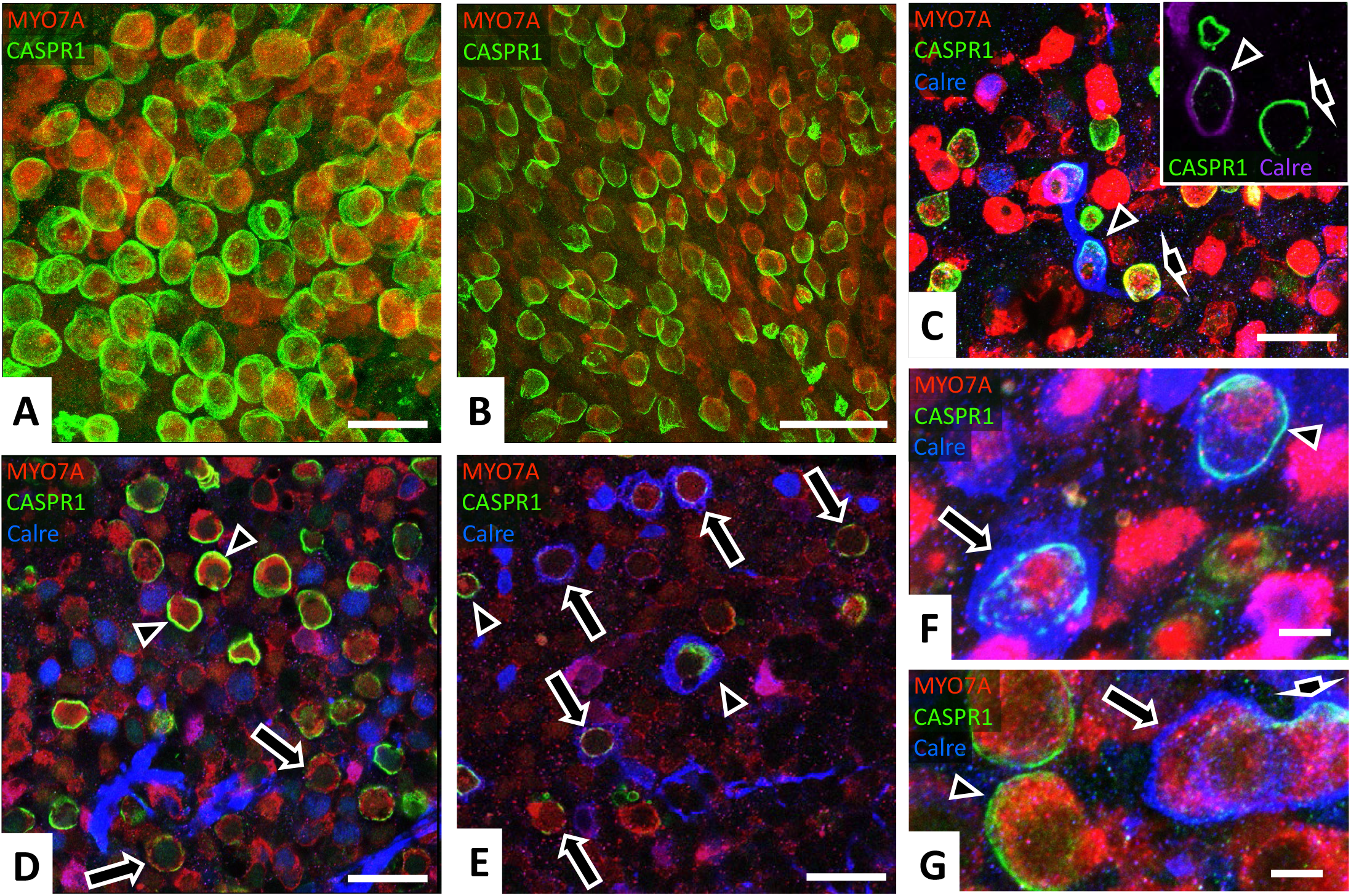
Evaluation of the calyceal junction in whole-mount vestibular epithelia from patients of vestibular diseases, as revealed by CASPR1 immunolabel (green, light blue when co-localizing with calretinin blue label). In the main panels, HC bodies are labelled with an anti-MYO7A antibody (red). Calretinin is shown in blue in main panels **C** to **F,** and in purple in the inset in **C. A.** Epithelium from a patient of Ménière’s Disease showing no pathological features. Note the elevated density of HCs and the continuous distribution of the CASPR1 label throughout the calyceal junction area of each HCI. B. Epithelium from a meningioma patient, showing also no overt pathology. **C.** Epithelium from a meningioma patient. Note the low density of HCs and the normally continuous CASPR1 label in calyces of both calretinin+ calyx-only terminals (arrowhead) and calretinin-dimorphic terminals (short arrow). The **inset** displays the same two calyces; a thinner stack is shown to highlight the uninterrupted CASPR1 label and its placement in the inner membrane of the calretinin+ calyx of the calyx-only terminal. D. Utricle periphery from a vestibular schwannoma patient. Around different HCI, the calyceal CASPR1 label is found in both normal continuous distribution (arrowhead) and in faint and broken distribution (long arrows). E. Striola region of the same utricle in D. Although some calyceal junctions display a roughly normal continuous distribution of the CASPR1 label (arrowheads), most display faint, broken or largely absent label (arrows). F. Higher magnification image from the same patient in D and E. Two calyces formed by calyx-only terminals are highlighted, one with continuous CASPR1 label (arrowhead) and the other with a patched distribution of this label (long arrow). G. Epithelium from a different vestibular schwannoma patient displaying normal CASPR1 distribution in calyces from dimorphic terminals (arrowhead) and a double calyx of a calretinin+ calyx-only terminal displaying a partial CASPR1 label in one HC (short arrow) and no CASPR1 in the other HC (long arrow). **Image thickness: A,** 12 μm; **B,** 10 μm; **C,** main 9.5 μm, inset 5 μm; **D,** 1.6 μm; **E,** 2.6 μm; **F,** 5 μm; **G,** 6 μm. **Scale Bars: A, C, D,** and **E,** 25 μm; **B,** 50 μm; **F,** and **G,** 5 μm.

## DISCUSSION

Previous work from our laboratory in rats and mice has demonstrated that sub-chronic IDPN exposure causes vestibular HC extrusion and that the extrusion is preceded by reversible calyceal junction dismantlement and synaptic uncoupling (Seoane et al., 2001a; Sedó-Cabezón et al., 2015; Greguske et al., 2019). These phenomena have received little research attention despite its potential relevance. HC extrusion is the main form of HC demise in birds and amphibians (Hirose et al., 1999; Rubel et al., 2013) and has been sporadically observed in mammalian vestibular epithelia (Li et al., 1995; Nakagawa et al., 1997; Granados and Meza, 2005). In the present work, we first aimed at establishing a new rat model of sub-chronic aminoglycoside toxicity to determine whether HC detachment and extrusion occur in ototoxic conditions of clinical relevance. The data obtained demonstrate that the calyceal junction reversibly dismantles during subchronic streptomycin exposure in association with reversible vestibular dysfunction. Synaptic uncoupling and HC extrusion were also recorded in these rats. In addition, we also collected evidence indicating that calyceal junction dismantlement may take place in the human vestibular system.

HC degeneration is effectively achieved by direct exposure to aminoglycoside antibiotics as done by trans-tympanic administration or in *in vitro* studies (Kohonen and Tarkkanen, 1969; Anniko et al., 1982). However, the use of more natural *in vivo* approaches to induce inner ear toxicity in laboratory animals with aminoglycosides has been a challenging issue, as acute systemic intoxication may not cause ototoxic effects until the dose is large enough to trigger animal death (Wu et al., 2001; Murillo-Cuesta et al., 2010). Nevertheless, several studies have succeeded in establishing dosing regimes that elicit ototoxic effects with limited lethality and systemic toxicity. Yet, many of these studies favour cochlear versus vestibular toxicity (Sullivan et al., 1987) or are designed to accelerate the progression of the damage (Taylor et al., 2008; Oesterle et al., 2008). In the present study, we successfully caused slowly progressing vestibular toxicity with streptomycin in young rats. By means of one daily injection, significant loss of vestibular function occurred after four weeks of the highest tolerated dose, while the use of two injections per day accelerated this loss in a dose-dependent manner, allowing for the study of ototoxicity emerging after 2-3 weeks of treatment. Interestingly, the effects were similar in both sexes and low variability was recorded within groups. Thus, as similarly reported for cochlear kanamycin toxicity in rats and mice (Wu et al., 2001; Jiang et al., 2006), exposure of Long-Evans rats to one or two daily doses of streptomycin reliably causes vestibular toxicity. Comparing the two models, it appears that the one daily dose regimen reaches a limit in its effect, while the two-dose regimen seems to provide a more flexible model to study dose- and time-dependent effects.

Streptomycin exposure in rats caused the loss of HCs, deeper after two daily doses than after one dose, as revealed by the SEM studies and quantified in immunofluorescence studies. Separate counts of HCI and HCII revealed that HC loss concentrated on the HCI, while HCII were spared in both injection paradigms. To identify HCI we used two different markers, CASPR1 and SPP1. CASPR1 is not expressed by HCIs but is found in the membrane of the calyx afferent contacting the HCI (Sousa et al., 2009; Lysakowski et al., 2011; Sedó-Cabezón et al., 2015). Although streptomycin caused a specific loss of CASPR1, traces of CASPR1 label in contact with cell profiles positive for Myo7a+ and negative for calretinin (see Fig 3A) allowed a reliable identification of HCI in the one dose samples. Also, except for a minor recuperation in HCI numbers in the periphery of utricles and cristae, no increase in HCI numbers was found in the samples of the recovery animals, in which a good recuperation in CASPR1 expression had occurred. In addition, the loss of HCI was confirmed using a positive HCI marker, SPP1 (McInturff et al., 2018), to evaluate the samples from the two daily injections experiment. To identify HCII, we also used a positive marker, calretinin, and found that the number of HCII per image was not reduced in any of the two exposure paradigms. Because of the method used, we cannot completely exclude that a small loss of HCII was masked by tissue shrinkage, but in any case the lack of differences in HCII numbers per image contrasts with the significant loss of HCI. While a greater sensitivity of HCI with respect to HCII in regards to ototoxic insult is widely recognized, not many studies are available in providing a quantitative assessment of this difference *in vivo* (Lopez et al., 1997; Nakayama et al., 1996; Hirvonen et al., 2005; Maroto et al., 2021a).

The data collected add to the existing evidence that HC extrusion is a significant form of HC demise induced by aminoglycosides in the mammalian vestibular system *in vivo*, as reported in the vestibular epithelia of rodents exposed to other aminoglycosides using relatively long exposure paradigms (Li et al., 1995; Granados and Meza, 2005). Also, the present data demonstrate that calyceal junction dismantlement and synaptic uncoupling occur after streptomycin, as previously demonstrated for sub-chronic IDPN (Sedó-Cabezón et al., 2015; Greguske et al., 2019). The streptomycin effect includes CASPR1 down-regulation, redistribution of KCNQ4, loss of electron density in the calyceal junction area, and a reduction in the number of synaptic puncta per cell. As observed in the IDPN model, these effects are associated with the loss of vestibular function, and significant recuperation was also recorded for the behavioural, CASPR1, and KCNQ4 effects in rats when allowed a recovery period after the exposure. However, significant differences between the two models were noticed. In the rat IDPN model (Seoane et al., 2001a; Sedó-Cabezón et al., 2015), a complete dismantlement of the calyceal junctions, with extensive loss of CASPR1 and redistribution of KCNQ4 occurs before HC loss starts. Extending the time of IDPN exposure triggers a massive event of HC extrusion, and the occurrence of extruding cells over several weeks indicates that the extrusion of each HC takes place over several days. IDPN thus provides a highly synchronic and slowly progressing model of sequential HC detachment and extrusion. In contrast, in the streptomycin rats, extruding HC were only found occasionally, the loss of CASPR1 and the redistribution of KCNQ4 were smaller, and a significant loss of HCs had occurred already before complete dismantlement of the calyceal junctions. These observations indicate that, in comparison to those of IDPN, the effects of streptomycin are less synchronic, and that the extrusion of each HC takes place in a shorter time, probably a few hours. Although not specifically addressed in the present experiments, the TEM studies did not give evidence for an extensive occurrence of apoptotic HC death. The simplest explanation of the present observations is that HC detachment and extrusion are triggered at the individual cell level, and that cells are not synchronized after streptomycin treatment as they are after IDPN. This conclusion is supported by our previous demonstration that calyceal junction dismantlement is triggered individually in a calyx-to-calyx basis also after IDPN (Sedó-Cabezón et al., 2015).

Behavioural and CASPR1 data presented here demonstrate repair capacity and functional recuperation of the rat vestibular epithelium after sub-chronic streptomycin exposure. Although only a limited number of animals were allowed to survive, their recuperation was indubitable. Functional recuperation may be a consequence of behavioural compensation or reflect recuperation from alterations not detected by histological analyses, such as hypothetical changes in endolymph composition or inactivation of the transduction channels. Conversely, our data excluded the possibility that HC regeneration accounted for this recuperation, because HCI counts remained decreased at the end of the recovery period. Rather, its association with the reversal in CASPR1 labelling, as measured on a cell-by-cell basis, suggests that the functional recuperation may result, at least in part, from repair phenomena in the HC and the contact between HCs and their afferents. The present observation of a recuperation in CASPR1 labelling contradicts the persistence of calyceal pathology observed in chinchillas exposed to gentamicin by direct perilymphatic infusion (Sultemeier and Hoffman, 2017). Differences in species, aminoglycosides, and exposure model may account for the difference between the present data and those by Sultemeier and Hoffman (2017).

To explore the possible relevance of the calyceal junction dismantlement to human pathology, we examined a series of human samples for histological evidence of such dismantlement. The interpretation of the human observations is weakened by the lack of control samples, and the variability in pathological conditions and surgery-generated damages. Nevertheless, many samples included a uniform CASPR1 label in all observable calyces, while some other samples offered multiple images of patched or largely missing label for this protein. Although the possibility that this abnormality was generated during collection of the sample cannot be completely excluded, the data strongly suggest that calyceal junction dismantlement may indeed occur in the human vestibule under stressful conditions. Although all the cases in which CASPR1 loss was recorded were from vestibular schwannoma surgeries, the number of samples from other pathologies was so low that it cannot be discarded that similar observations could emerge if more cases were examined. Current evidence indicates that vestibular schwannomas may damage the sensory epithelia of the inner ear through tumour-secreted factors, including tumour necrosis factor alpha (TNFα) (Dilwali et al., 2015). A recent study has demonstrated the potential of this pro-inflammatory cytokine to cause cochlear synaptic uncoupling *in vivo* (Katsumi et al., 2020). It is thus possible that inflammatory signals like TNFα are able to trigger the dismantlement of the calyceal junction, synaptic uncoupling, and eventual HC extrusion in the vestibular epithelium.

Future work may address some of the limitations of this initiating study and use it to expand our understanding of vestibular pathophysiology. For instance, determining the blood and inner ear concentrations of streptomycin (and its potential ototoxic metabolites, e.g. Granados and Meza, 2005) in this exposure model may enlighten the role of inner ear accumulation in systemic aminoglycoside toxicity. Recent data supports that cochlear retention is a key factor in cisplatin ototoxicity (Breglio et al., 2017), and some data is available supporting a similar conclusion for the cochlear and vestibular toxicity of gentamicin (Imamura and Adams, 2003). More works is also needed to replicate and expand the human observations, excluding any potential confounding factor and relating the pathological findings to the clinical data of patients. This is particularly important due to the lack of tissues from humans not suffering any inner ear disease.

In conclusion, the present study evaluated whether events leading to HC extrusion in the vestibular sensory epithelia of rodents exposed to sub-chronic IDPN (Seoane et al., 2001a; Sedó-Cabezón et al., 2015; Greguske et al., 2019) may have a wider significance. The data collected demonstrate that dismantlement of the calyceal junction and synaptic uncoupling are widespread phenomena in rats exposed to sub-chronic streptomycin regimes that cause HC loss through, at least in part, the extrusion pathway. We also collected data suggesting that calyceal junction dismantlement may occur in human vestibular epithelia in response to inflammatory stress. Elucidating the molecular pathways triggering these cellular phenomena may unveil new targets to control damage progression in this sensory epithelium.

## Supporting information

Supplemental Fig S8 Animated pptx

## Acknowledgements

This study was supported by grant number RTI2018-096452-B-I00 and PID2021-124678OB-I00 (Ministerio de Ciencia e Innovación, Agencia Estatal de Investigación, MCIU/AEI, 10.13039/501100011033, and European Regional Development Fund, FEDER), grant number 2017SGR621 (Agència de Gestió d’Ajuts Universitaris i de Recerca, AGAUR, Generalitat de Catalunya), and grant number 202007-30-31 (Fundació La Marató de TV3 2019). A.B.G is a Serra-Húnter fellow. The electron and confocal microscopy studies were performed at the Centres Científics i Tecnològics de la UB (CCiTUB) of the Universitat de Barcelona. We thank Benjamin Torrejon-Escribano, Rosa Rivera, Adriana Martínez Gené, Mora Gimeno, Josep Manel Rebled and Eva Prats for the expert and technical help. We also thank Dr Bechara Kachar for the generous gift of the KCNQ4 antibody and Dr. Erin A. Greguske for revision of the manuscript. We are also grateful to the patients that gave consent to the research use of their sensory tissues.

