## Supplemental Tables for "The vestibular calyceal junction is dismantled following subchronic streptomycin in rats and sensory epithelium stress in humans"

**SUPPLEMENTARY TABLE S1**

Animals and treatments. Streptomycin was given s.c., 1 dose/day or 2 doses/day.

| **Experiment number** | **Daily dose (mg/kg)** | **Time (weeks)** | **Male** | **Female** | | **At end of exposure** | **Recovery** |
|---|---|---|---|---|---|---|---|
| E0526 | 0 | 6 | 4 | 0 | | 4 | 0 |
| | 100 | 6 | 4 | 0 | | 4 | 0 |
| | 300 | 6 | 3 | 0 | | 3 | 0 |
| | 300 | died | 1 | 0 | | - | - |
| | 500 | 6 | 3 | 0 | | 3 | 0 |
| | 500 | died | 1 | 0 | | - | - |
| E0531 | 0 | 6 or 8 | 0 | 4 | | 2 | 2 |
| | 400 | 8 | 0 | 4 | | 2 | 2 |
| | 500 | 6 | 0 | 3 | | 3 | 0 |
| | 500 | 8 | 0 | 5 | | 3 | 2 |
| E0532 | 0 | 6 or 8 | 6 | 0 | | 4 | 2 |
| | 500 | 6 | 3 | 0 | | 0 | 3 |
| | 500 | 8 | 7 | 0 | | 4 | 3 |
| | 500 | died | 2 | 0 | | - | - |
| E0538 | 0 | 6 | 4 | 4 | | 8 | 0 |
| | 500 | 6 | 4 | 4 | | 8 | 0 |
| E0545 | 0 | 4 | 1 | 1 | | 2 | 0 |
| | 0 | 6 | 3 | 3 | | 6 | 0 |
| | 600 (2 x 300) | 3 | 1 | 0 | | 1 | 0 |
| | 600 (2 x 300) | 6 | 3 | 4 | | 8 | 0 |
| | 700 (2 x 350) | 3 | 0 | 1 | | 0 | 1 |
| | 700 (2 x 350) | 4 | 4 | 3 | | 6 | 1 |
| | 800 (2 x 400) | 3 | 4 | 4 | | 6 | 2 |
| **TOTAL** | | | **58** | **40** | | | |
| **Pooled experiments** | **Daily dose (mg/kg)** | **Time (weeks)** | **Male** | **Female** | | **At end of exposure** | **Recovery** |
| | 0 | 4-6-8 | 18 | 12 | | 26 | 4 |
| | 100 | 6 | 4 | 0 | | 4 | 0 |
| | 300 | 6 | 3 | 0 | | 3 | 0 |
| | 400 | 8 | 0 | 4 | | 2 | 2 |
| | 500 | 6 | 10 | 7 | | 14 | 3 |
| | 500 | 8 | 7 | 5 | | 7 | 5 |
| | 600 (2 x 300) | 3 | 1 | 0 | | 0 | 1 |
| | 600 (2 x 300) | 6 | 3 | 4 | | 8 | 0 |
| | 700 (2 x 350) | 3 | 0 | 1 | | 0 | 1 |
| | 700 (2 x 350) | 4 | 4 | 3 | | 6 | 1 |
| | 800 (2 x 400) | 3 | 4 | 4 | | 6 | 2 |
| | 300 | died | 1 | 0 | | - | - |
| | 500 | died | 3 | 0 | | - | - |
| **TOTAL** | | | **58** | **40** | | | |

**SUPPLEMENTARY TABLE S2**

Animals used in histological analyses. Each animal in the study was used in several analyses.

**Scanning Electron Microscopy**

| Dose (mg/kg·day) | Time (weeks) | At end of exposure | Recovery |
|---|---|---|---|
| 0 | 4, 6 or 8 | 12 | 4 |
| 300 | 6 | 3 | 0 |
| 400 | 8 | 2 | 2 |
| 500 | 6 | 6 | 3 |
| 500 | 8 | 7 | 5 |
| 600 (2 x 300) | 6 | 7 | 0 |
| 700 (2 x 350) | 4 | 6 | 2 |
| 800 (2 x 400) | 3 | 6 | 2 |

**Transmission Electron Microscopy**

| Dose (mg/kg·day) | Time (weeks) | At end of exposure |
|---|---|---|
| 0 | 4, 6 or 8 | 4 |
| 500 | 6 | 5 |
| 500 | 8 | 4 |
| 600 (2 x 300) | 6 | 3 |
| 700 (2 x 350) | 4 | 2 |

**Immunohistochemistry**

| Dose group (mg/kg·day) | Time (weeks) | I. 1 | I. 2 | I. 3 | I. 4 | I. 5 |
|---|---|---|---|---|---|---|
| 0 | 4-6-8 | 10 | 8 | 5 | 5 | 4 |
| 500 | 6 | 6 | - | 3 | 4 | - |
| 500 | 8 | 7 | - | 5 | - | - |
| 500-Washout | 6-8 | 8 | - | 5 | - | - |
| 600 (2 x 300) | 6 | - | 7 | - | - | 7 |
| 700 (2 x 350) | 4 | - | 6 | - | - | 5 |
| 800 (2 x 400) | 3 | - | 6 | - | - | 6 |
| 700/800 (2x350/400) Washout | 3-4 | - | 4 | - | - | - |

**I.1:** Immunohistochemical analysis number 1, data in Figures 3 and 5. HCI and HCII counts and CASPR1 expression, one injection per day, labelling with antibodies against MYO7A, CASPR1 and Calretinin.
**I.2:** Data in Figure 4 and Supplementary Figure S5. HCI and HCII counts and CASPR1 expression, two injections per day, labelling with antibodies against SPP1, CASPR1, Calretinin.
**I.3:** Data in Figure 6. KCNQ4 expression, one injection per day after labelling with antibodies against CASPR1 and KCNQ4.
**I.4:** Data in Figure 7. Synaptic puncta counts, one injection per day, labelling with antibodies against MYO7A, CASPR1, Ribeye and PSD-95.
**I.5:** Data in Figure 9. Observation of HC extrusion, two injections per day, labelling with antibodies against MYO7A, CASPR1, Radixin.

**SUPPLEMENTARY TABLE S3**

Pathology, age and sex of the patients included in the study. Samples from patients with the following pathologies were examined: Ménière's disease (MD, n=2), vestibular schwannoma (VS, n=22), meningioma (Men., n=4), and endolymphatic sac tumour (ELST, n=1). The right column shows the time (in days) between surgery and delivery of the sample to the laboratory, during which the tissue remained in fixative. For several months, the time of delivery was not recorded (N.R.), but was similar to that in other periods (i.e., most samples were received the day after surgery.

| Sample | Pathology | Age | Sex | Fixation |
|---|---|---|---|---|
| 3 | MD | 58 | F | 1 |
| 8 | VS | 60 | M | 1 |
| 9 | VS | 51 | M | 1 |
| 10 | VS | 51 | M | 1 |
| 11 | VS | 46 | M | 4 |
| 14 | Men. | 68 | F | 1 |
| 15 | VS | 63 | F | 1 |
| 16 | MD | 58 | F | 1 |
| 17 | VS | 18 | M | 1 |
| 18 | VS | 45 | M | 1 |
| 19 | Men. | 47 | F | 1 |
| 21 | VS | 50 | M | 1 |
| 22 | VS | 54 | M | 4 |
| 25 | VS | 67 | M | 1 |
| 31 | VS | 62 | M | N.R. |
| 32 | VS | 66 | F | N.R. |
| 33 | VS | 31 | F | N.R. |
| 35 | VS | 53 | M | N.R. |
| 36 | VS | 56 | M | N.R. |
| 41 | VS | 36 | M | N.R. |
| 42 | VS | 48 | F | N.R. |
| 43 | VS | 59 | F | N.R. |
| 45 | VS | 53 | F | N.R. |
| 52 | ELST | 69 | F | N.R. |
| 57 | Men. | 57 | F | 1 |
| 59 | VS | 37 | M | 7 |
| 60 | VS | 55 | F | 1 |
| 64 | VS | 70 | F | 4 |
| 89 | Men. | 41 | F | 1 |

**SUPPLEMENTARY TABLE S4 - Antibodies**

| Target | Host and type | Source | Specificity |
|---|---|---|---|
| Caspr1 | Mouse monoclonal (IgG1) | Clone K65/35, Neuromab, RRID: AB_2083496 | Western blot on rat brain membranes from control and caspr1-KO mice (Datasheet). |
| PSD-95 | Mouse monoclonal (IgG2a) | Clone K28/43, Neuromab, RRID: AB_2292909 | Western blot on mouse brain membranes from control and PSD-95-KO mice (Datasheet). |
| Ribeye | Mouse monoclonal (IgG1) | Clone 16/CtBP2, BD Biosciences, RRID: AB_399431 | Western blot of CtBP2 on cell lysates (Datasheet). See also Pujol et al., 2014 |
| Myosin VIIa | Rabbit polyclonal | 25-6790, Proteus Biosciences, RRID: AB_10015251 | Validated by immunoblot, widely used for specific hair cell labelling. See Pujol et al., 2014. |
| Calretinin | Guinea pig polyclonal | 214.104, Synaptic Systems, RRID: AB_10635160 | Co-localizes with anti-calretinin antibodies validated in KO mice (CR 7699 from Swant). |
| SPP1 (Ospeopontin) | Goat polyclonal | AF808, RD Systems, RRID: AB_2194992 | Specifically labels type I hair cells as described (McInturff et al., 2018), in perfect concordance with CASPR1 label of encasing calyces (Sousa et al., 2009). |
| KCNQ4 | Rabbit polyclonal | Donated by Bechara Kachar (PB180) | Beisel et al., 2005.<br>Label co-localized with other validated anti-KCNQ4 antibodies (Spitzmaul et al., 2013; see Sedó-Cabezón et al., 2015) |

Beisel, K.W., Rocha-Sanchez, S.M., Morris, K.A., Nie, L., Feng, F., Kachar, B., Yamoah, E.N., Fritzsch, B (2005) Differential expression of KCNQ4 in inner hair cells and sensory neurons is the basis of progressive high-frequency hearing loss. *J. Neurosci.* 25, 9285–9293.

Lysakowski, A., Gaboyard-Niay, S., Calin-Jageman, I., Chatlani, S., Price, S.D. and Eatock, R.A. (2011). Molecular microdomains in a sensory terminal, the vestibular calyx ending. *J. Neurosci*. 31, 10101–10114.

McInturff, S., Burns, J. C. and Kelley, M. W. (2018). Characterization of spatial and temporal development of Type I and Type II hair cells in the mouse utricle using new cell-type-specific markers. *Biol. open*, 7(11).

Pujol R, Pickett SB, Nguyen TB and Stone JS (2014) Large basolateral processes on type II hair cells are novel processing units in mammalian vestibular organs. *J. Comp. Neurol.* 522, 3141-3159. doi: 10.1002/cne.23625. 10.

Sedó-Cabezón, L., Jedynak, P., Boadas-Vaello, P. and Llorens, J. (2015) Transient alteration of the vestibular calyceal junction and synapse in response to chronic ototoxic insult in rats. *Dis. Model. Mech.* 8, 1323-1337.

Sousa, A.D., Andrade, L.R., Salles, F.T., Pillai, A.M., Buttermore, E.D., Bhat, M.A. and Kachar, B (2009). The septate junction protein caspr is required for structural support and retention of KCNQ4 at calyceal synapses of vestibular hair cells. *J. Neurosci*. 29, 3103-3108.

Spitzmaul, G., Tolosa, L., Winkelman, B.H., Heidenreich, M., Frens, M.A., Chabbert, C., de Zeeuw, C.I. and Jentsch, T.J. (2013) Vestibular role of KCNQ4 and KCNQ5 K+ channels revealed by mouse models. *J. Biol. Chem.* 288, 9334-9344.
