## Supplemental Figures for "The vestibular calyceal junction is dismantled following subchronic streptomycin in rats and sensory epithelium stress in humans"

SUPPLEMENTARY FIGURE S1

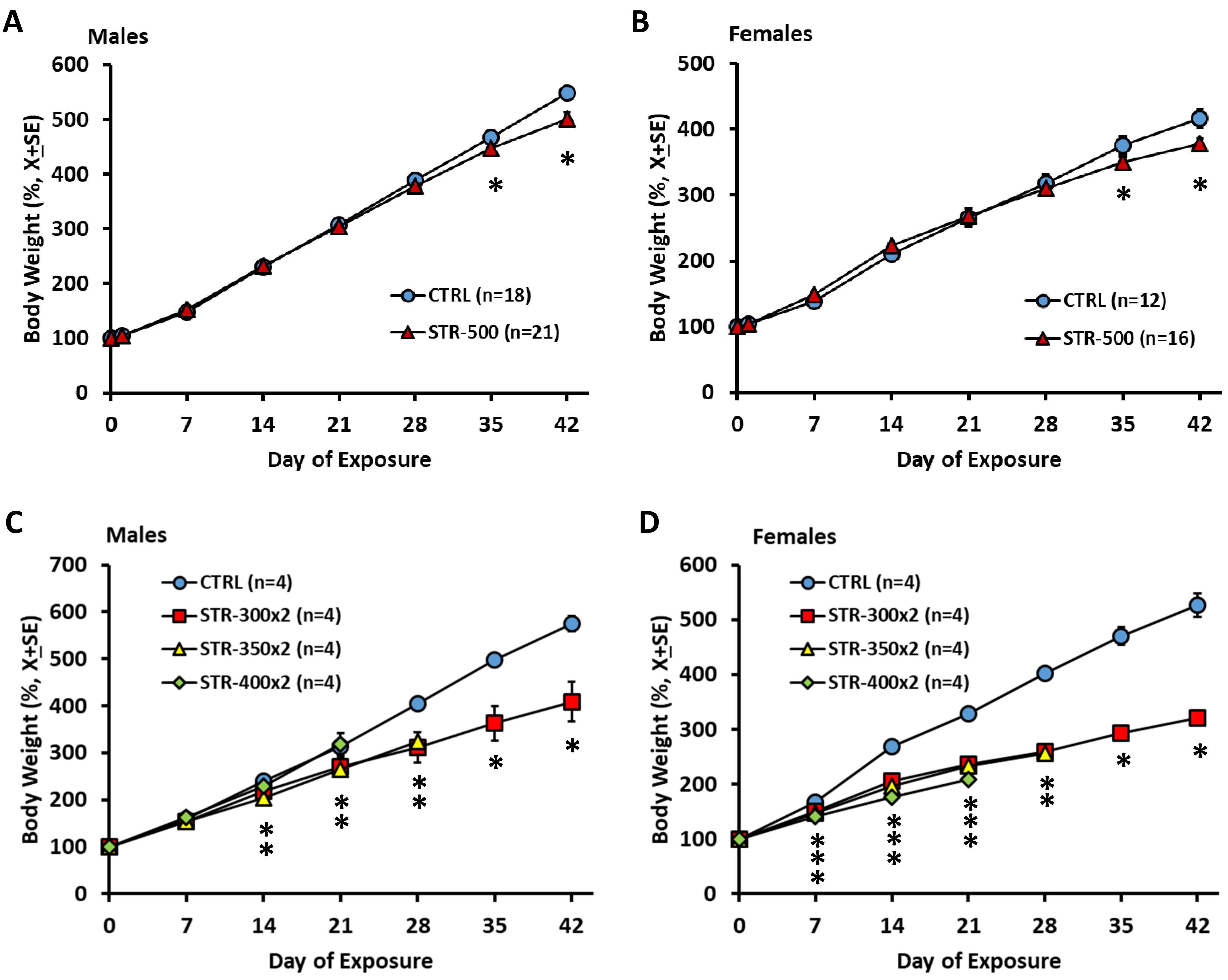


**Supplementary figure S1.** Effects of streptomycin on body weight in male and female Long-Evans rats. **A and B.** Effects of one s.c. injection per day at 0 (vehicle control, PBS) or 500 mg/kg (as free base) of streptomycin sulphate starting at 21 days of age, for 6 weeks. Data are mean body weights (+SE), in percentage of initial weights. Both control and treated animals showed a constant increase in body weight, but streptomycin reduced the rate of increase by the end of the exposure period. In males, repeated-measures MANOVA resulted in significant day ($F[7,31]=764.0$, $p<0.001$), treatment ($F[1,37]=5.48$, $p=0.025$), and day X treatment interaction ($F[7,31]=15.28$, $p<0.001$) effects. In females, the analysis resulted in significant day ($F[7,20]=372.6$, $p<0.001$) and day X treatment interaction ($F[7,20]=9.50$, $p<0.001$) effects, but no significant treatment effect ($F[1,26]=1.52$, $p=0.229$). **C and D.** Effects of two s.c. injections per day of 0 (vehicle control, PBS), 300, 350, or 400 mg/kg (as free base) per dose of streptomycin starting at 21 days of age, for 4-6, 6, 4, or 3 weeks, respectively. Data are mean body weights (+SE) in percentage of initial weights. In males, the numbers of animals were reduced to 3 in the CTRL group after day 28 and in the STR-300x2 after day 21. In females, the numbers of animals were reduced to 3 in the CTRL group after day 28 and in the STR-350x2 after day 21. Both control and treated animals showed a constant increase in body weight, but streptomycin reduced the rate of increase from the first (females) or second (males) weeks of treatment. In males, repeated-measures MANOVA of the data from days 7 to 21 resulted in significant day ($F[2,11]=353.8$, $p<0.001$), treatment ($F[3,12]=8.076$, $p=0.003$), and day X treatment interaction ($F[6,22]=4.698$, $p=0.003$) effects. In females, the analysis resulted in significant day ($F[2,11]=101.4$, $p<0.001$), treatment ($F[3,12]=7.684$, $p=0.004$), and day X treatment interaction ($F[6,22]=4.105$, $p=0.007$) effects.* $p<0.05$, different from control group, Dunnett's test after significant ANOVA at that day.

SUPPLEMENTARY FIGURE S2

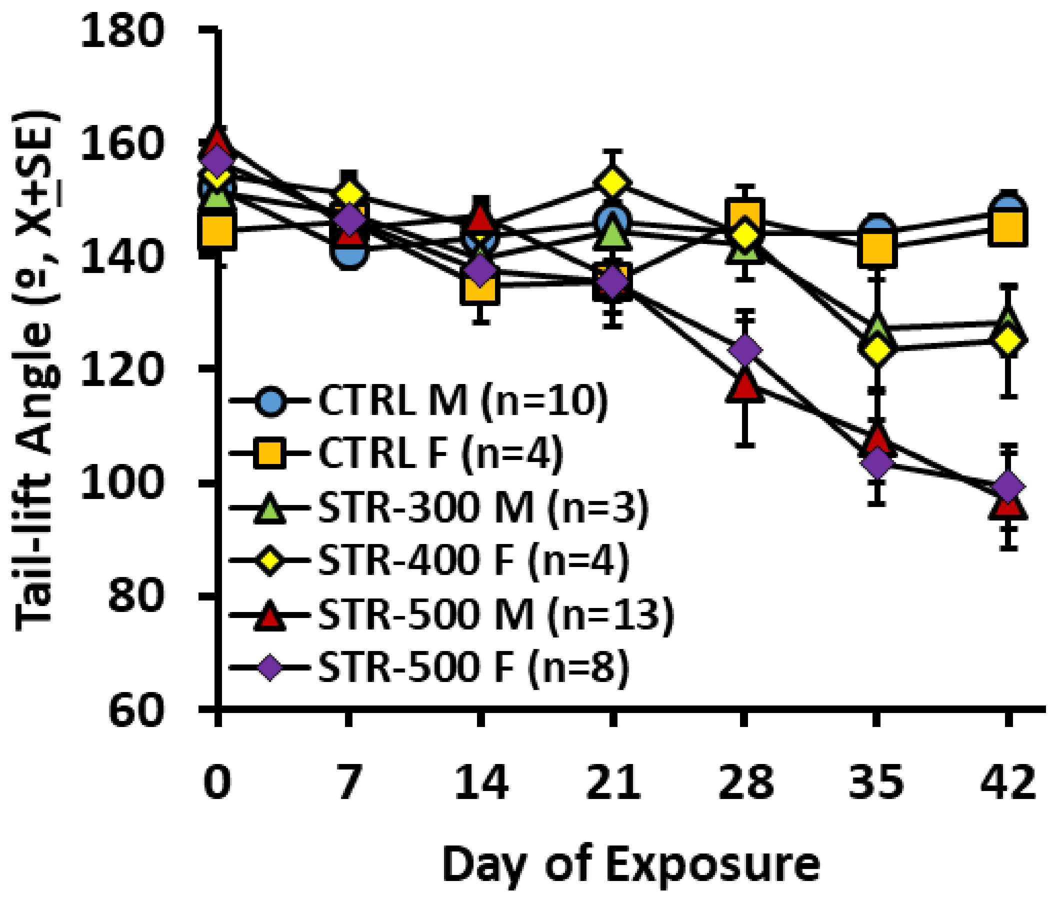


**Supplementary figure S2.** Effects of streptomycin on the tail-lift angle as a function of dose and sex. Male (M) and female (F) rats received one s.c. injection per day at 0 (vehicle control, PBS - CTRL), 300 (only males), 400 (only females), or 500 mg/kg of streptomycin sulphate (as free base) starting at 21 days of age, for 6 weeks. To evaluate the impact of sex on the effect, we performed a MANOVA analysis with control animals and animals administered with the 500 mg/kg dose. The analysis indicated a significant effect of the day factor ($F[6,26]=12.28$, $p<0.001$), the treatment factor ($F[1,31]=13.61$, $p<0.001$) and the day X treatment interaction ($F[6,26]=7.13$, $p<0.001$). Not significant were the sex factor ($F[1,31]=0.341$, $p=0.563$), the sex X day interaction ($F[6,26]=1.42$, $p=0.246$), the sex X treatment interaction ($F[1,31]=0.085$, $p=0.772$), and the sex X treatment X day interaction ($F[6,26]=0.261$, $p=0.950$).

SUPPLEMENTARY FIGURE S3

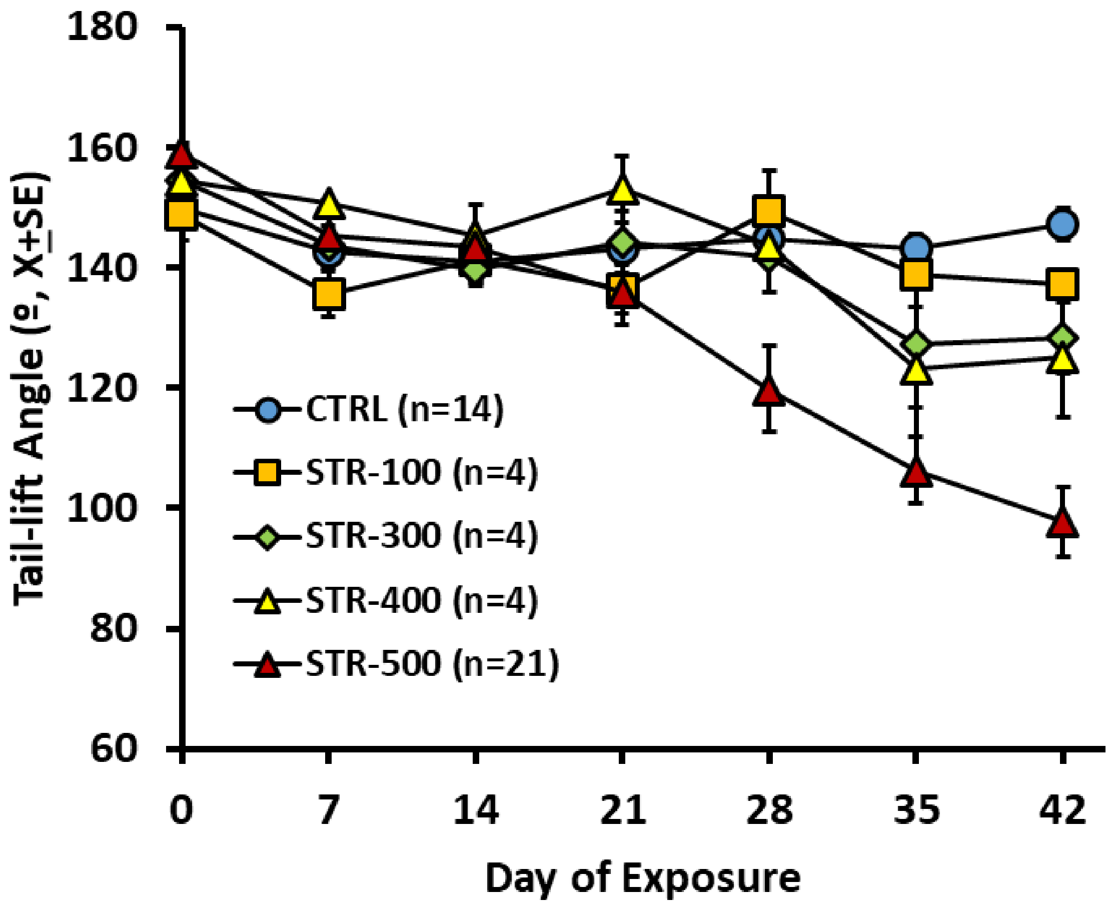


**Supplementary figure S3.** Effects of streptomycin on the tail-lift angle. Rats received one s.c. injection per day at 0 (vehicle control, PBS), 100, 300, 400, or 500 mg/kg (as free base) of streptomycin sulphate starting at 21 days of age, for 6 weeks.

SUPPLEMENTARY FIGURE S4

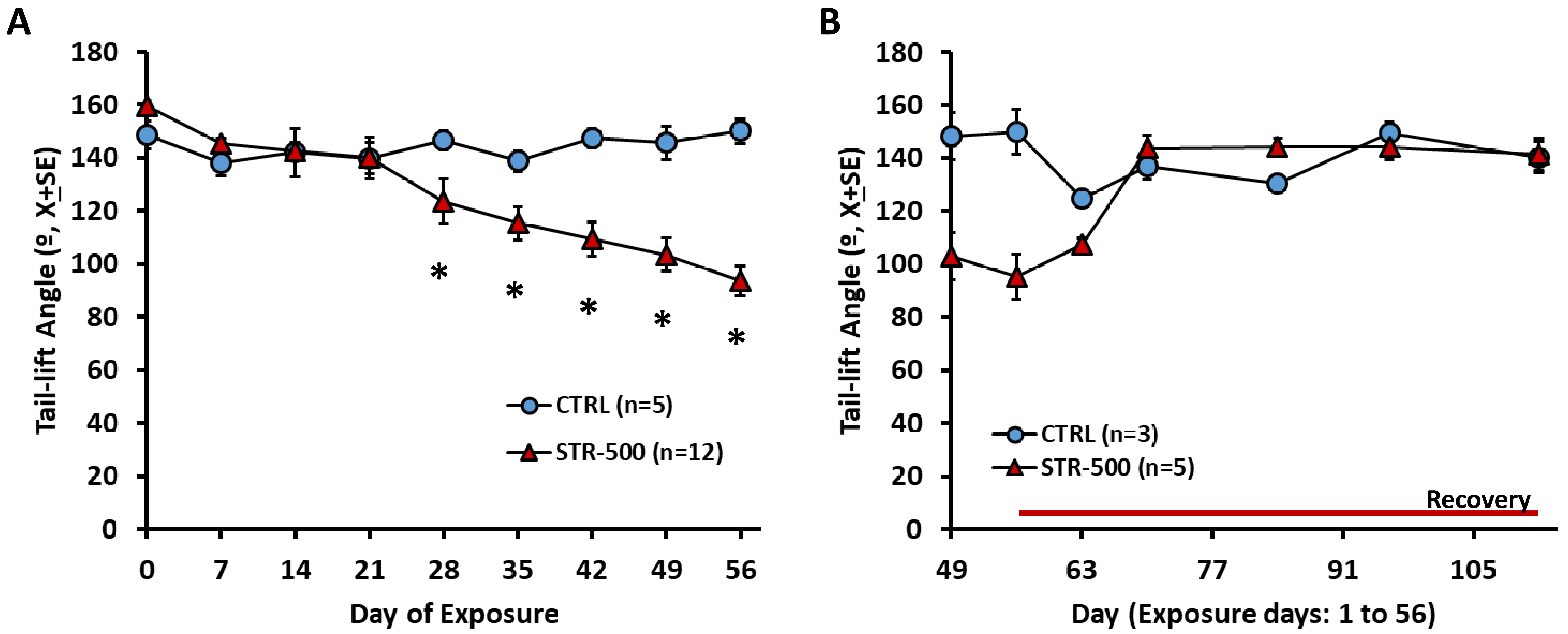


**Supplementary figure S4.** Effects of streptomycin on the tail-lift angle. **A.** Effects of one s.c. injection per day at 0 (vehicle control, PBS) or 500 mg/kg (as free base) of streptomycin sulphate starting at 21 days of age, for 8 weeks. Repeated-measures MANOVA resulted in significant day ($F[8,8]=7.067$, $p=0.006$), treatment ($F[1,15]=12.303$, $p=0.003$), and day X treatment interaction ($F[8,8]=5.044$, $p=0.017$) effects. * $p<0.05$, different from control group, day by day post-hoc analysis. **B.** Tail-lift angles after the exposure period in the animals allowed to survive. The data at 49 and 56 days are a subset of the data also shown in panel A. The animals included in A and not in B were processed for histological analysis at 8 weeks of exposure.

SUPPLEMENTARY FIGURE S5

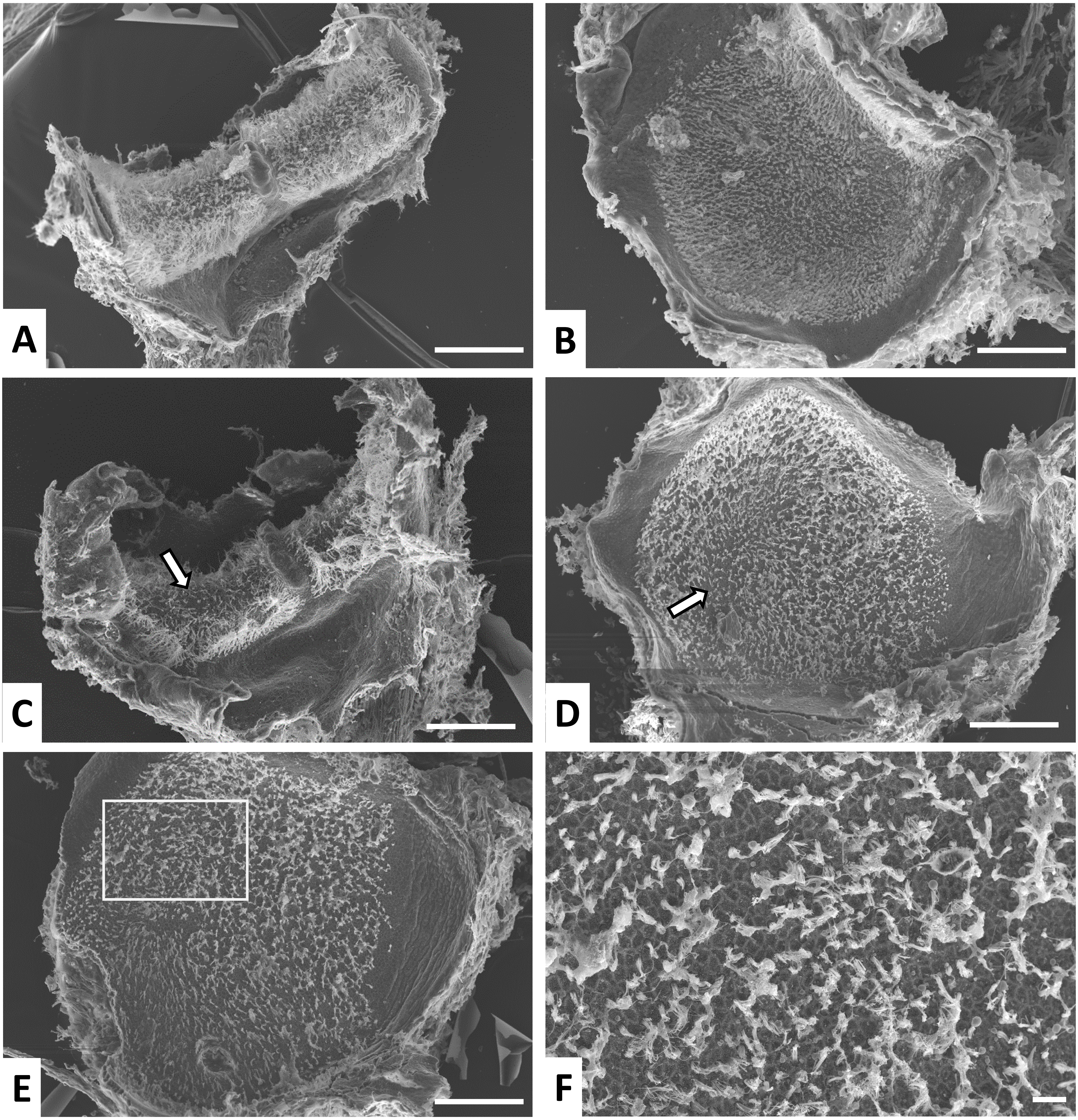


**Supplementary figure S5.** Effects of sub-chronic streptomycin (500 mg/kg, once a day, for 8 weeks) on the vestibular sensory epithelium of the rat as observed in surface preparations of cristae and utricles examined by SEM. **A and B.** Control crista and utricle, respectively. Note the high density of stereocilia bundles throughout the epithelium. **C and D.** Effect of streptomycin in samples representative of the average effect. A reduced density of stereocilia bundles is noticeable in the apical part of the crista and the striola region of the utricle (arrows). **E.** Worst-case example, with an overt reduction in the density of hair bundles throughout the epithelium. The boxed area is shown at higher magnification in panel **F.** The lack of HCs is revealed by the borders of the supporting cells, drawn by the high densities of microvilli at the cell borders. **Scale Bars:** 100 mm in **A, B, C , D,** and **E;** 10 mm in **F**

SUPPLEMENTARY FIGURE S6

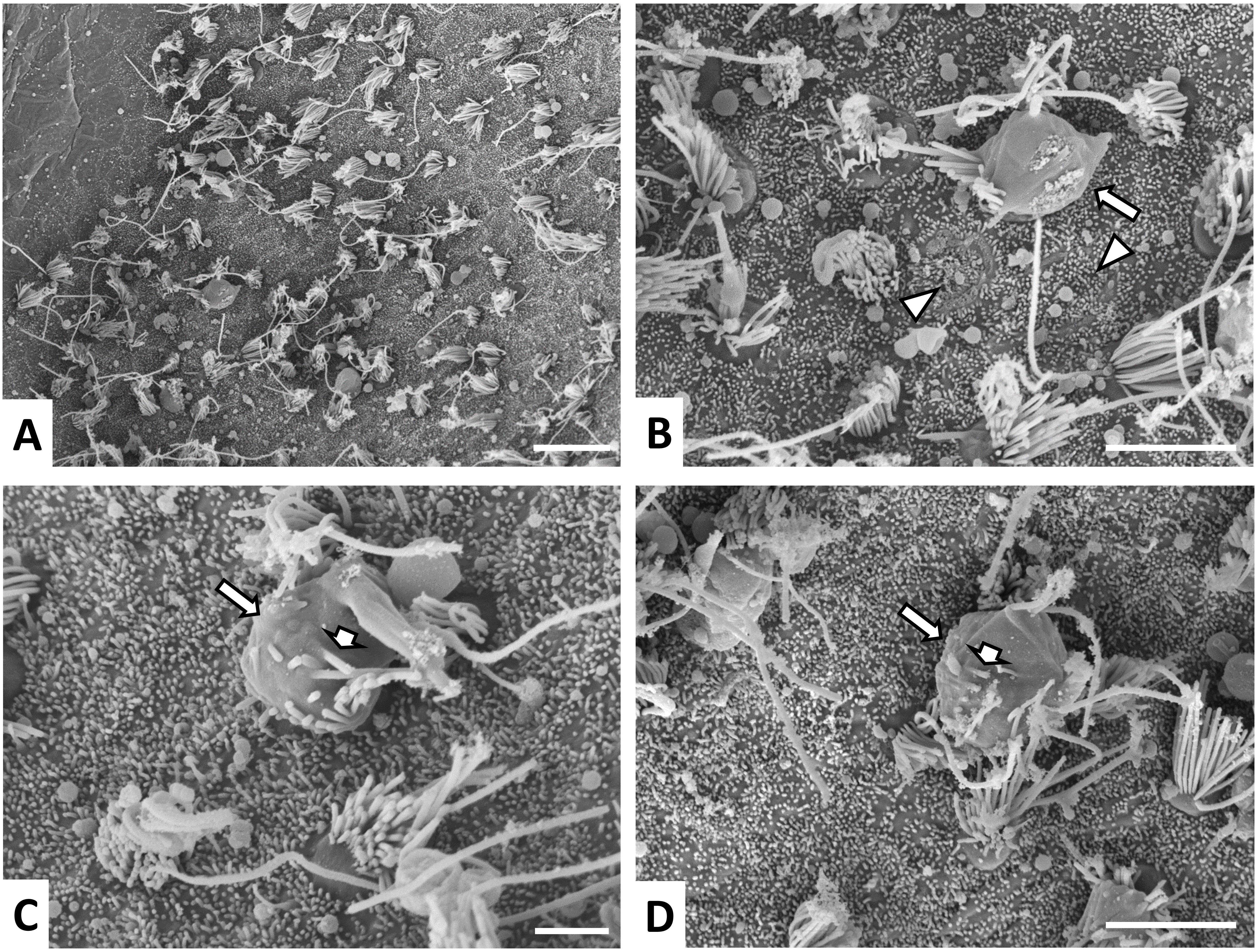


**Supplementary figure S6.** Effects of sub-chronic streptomycin (400 mg/kg, twice a day, for 3 weeks) on the vestibular sensory epithelium of the rat as observed in surface preparations of utricles examined by SEM. **A.** General view. Note the overtly reduced density of hair bundles, and the presence of protruding cells or fused stereocilia. **B-D**. Higher magnification images highlighting evidences of ongoing damage, including the presence of extruding hair cells (long arrows in the three panels) and of epithelial scars (arrowheads in B). Short arrows in **C** and **D** point to stereocilia that allow to recognize that the extruding cell is a HC. **Scale Bars:** 10 mm in **A;** 5 mm in **B** and **D;** 2 mm in **C.**

SUPPLEMENTARY FIGURE S7

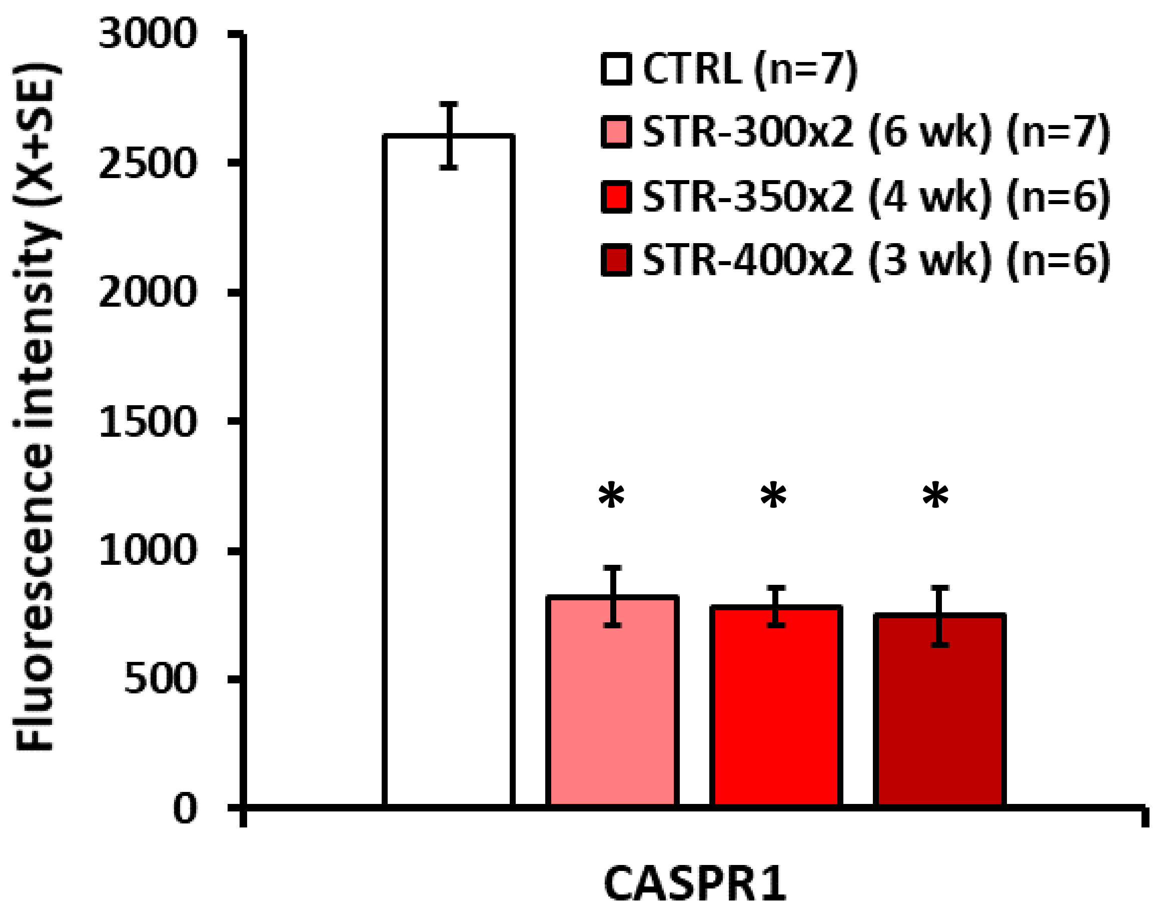


**Supplementary figure S7.** Effects of sub-chronic streptomycin on the expression of CASPR1 in the vestibular crista of the rat. Animals received two injections per day at 0 (CTRL, vehicle control), 300 (STR-300x2), 350 (STR-350x2), or 400 (STR-400x2) mg/kg of streptomycin for 6, 6, 4, or 3 weeks, respectively, starting at 21 days of age. Data are fluorescence amount per cell +/- SE in arbitrary units. A single value was obtained from each animal by averaging the fluorescence intensity in all available HCI in 6 (in a few cases, 5 or 7) sections of a crista. N = number of animals per group. *: different from CTRL ($p<0.05$), Duncan's test after significant ANOVA ($F[3,22]=74.91$, $p<0.001$.

SUPPLEMENTARY FIGURE S8

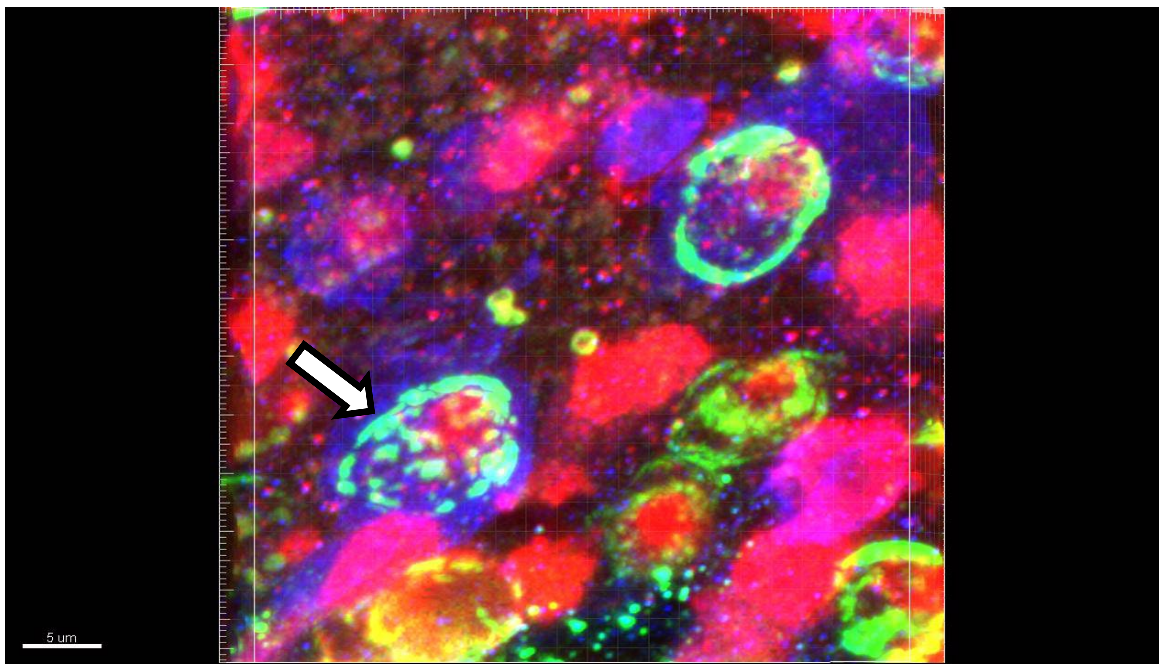


**Supplementary figure S8.** Animated 3D reconstruction of a vestibular epithelium from a vestibular schwannoma patient. The epithelium was immunostained with anti-MYO7a (red), anti-calretinin (blue), and anti-CASPR1 (green) antibodies. The apparent co-localisation of calretinin and CASPR1 (light blue), corresponds to the calyceal junction (CASPR1+) of a calretinin+ calyx-only terminal. Note the patched appearance of the calyceal junction marked with an arrow at the beginning of the video.

To view the animation, open the accompanying ppt file and activate it.
