## Supplemental Fig S8 Animated pptx for "The vestibular calyceal junction is dismantled following subchronic streptomycin in rats and sensory epithelium stress in humans"

### Slide 1
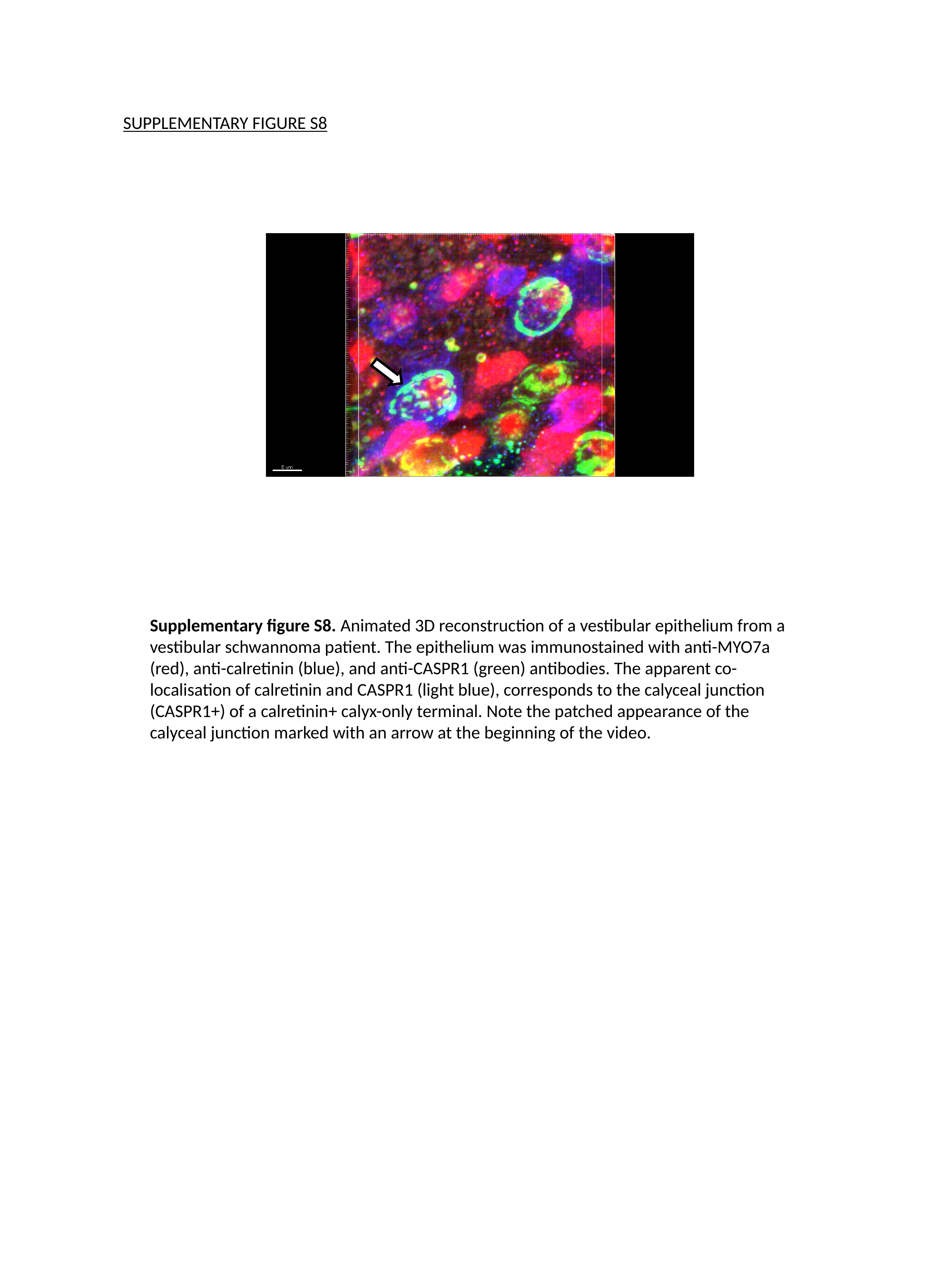

SUPPLEMENTARY FIGURE S8
Supplementary figure S8. Animated 3D reconstruction of a vestibular epithelium from a vestibular schwannoma patient. The epithelium was immunostained with anti-MYO7a (red), anti-calretinin (blue), and anti-CASPR1 (green) antibodies. The apparent co-localisation of calretinin and CASPR1 (light blue), corresponds to the calyceal junction (CASPR1+) of a calretinin+ calyx-only terminal. Note the patched appearance of the calyceal junction marked with an arrow at the beginning of the video.
